# Molecular mechanism of a parasite kinesin motor and implications for its inhibition

**DOI:** 10.1101/2021.01.26.428220

**Authors:** Alexander D. Cook, Anthony Roberts, Joseph Atherton, Rita Tewari, Maya Topf, Carolyn A. Moores

## Abstract

*Plasmodium* parasites cause malaria and are responsible annually for hundreds of thousands of deaths. They have a complex life cycle in which distinct stages are transmitted between, and reproduce in, human and mosquito hosts. In the light of emerging resistance to current therapies, components of the parasite replicative machinery are potentially important targets for anti-parasite drugs. Members of the superfamily of kinesin motors play important roles in the microtubule-based replicative spindle machinery, and kinesin-5 motors are established anti-mitotic targets in other disease contexts. We therefore studied kinesin-5 from *Plasmodium falciparum* (*Pf*K5) and characterised the biochemical properties and structure of the *Pf*K5 motor domain. We found that the *Pf*K5 motor domain is an ATPase with microtubule plus-end directed motility. We used cryo-EM to determine the motor’s microtubule-bound structure in no nucleotide and AMPPNP-bound states. Despite significant sequence divergence in this motor, these structures reveal that this parasite motor exhibits classical kinesin mechanochemistry. This includes ATP-induced neck-linker docking to the motor domain, which is consistent with the motor’s plus-ended directed motility. Crucially, we also observed that a large insertion in loop5 of the *Pf*K5 motor domain creates a dramatically different chemical environment in the well characterised human kinesin-5 drug-binding site. Our data thereby reveal the possibility for selective inhibition of *Pf*K5 and can be used to inform future exploration of *Plasmodium* kinesins as anti-parasite targets.

## Introduction

Malaria is a massive disease burden world-wide, with an estimated 219 million cases in 2017, a year which also saw the first increase in cases for nearly two decades ^1^. With resistance to current frontline therapeutics rapidly rising ^2–4^, new drug targets are urgently needed. Malaria is caused by *Plasmodium* parasites, which are unicellular eukaryotes belonging to the Apicomplexa phylum. Malaria parasites have a complex life-cycle involving distinct stages that are transmitted between, and reproduce in, human and mosquito hosts ^5^. The cytoskeleton plays an important role throughout the parasite life cycle, and the microtubule (MT) based spindle machinery is involved in the many rounds of mitotic and meiotic replication required for parasite proliferation. Anti-mitotics are well-established as drugs in a variety of settings, notably human cancer ^6^ – thus, components of the malaria replicative machinery are attractive anti-parasite targets. However, given the obligate intracellular nature of malaria parasites, any therapeutic target must be sufficiently divergent to be selectively disrupted compared to host homologues.

Members of the kinesin superfamily are such potential targets. Kinesins are motor proteins that bind to MTs and convert the energy of ATP binding and hydrolysis into MT-based mechanical work. Different kinesin families have specialised functions, such as translocation of cargo along MTs, regulation of MT polymer dynamics, and organisation of higher-order MT structures like mitotic and meiotic spindles ^7,8^. MTs are built from heterodimers of the highly conserved α- and β-tubulin and, whereas there is approximately 95% sequence conservation of tubulins between *Plasmodium sp*. and *Homo sapiens*, sequence conservation within kinesin families is much lower, typically 40-50 %. This raises two important questions: do distantly related members of kinesin families diverge in their molecular properties, and could such sequence divergence allow selective inhibitors to be developed?

The kinesin-5 family are involved in cell division in many organisms, have long been investigated as an anti-mitotic therapeutic target for human cancer ^9^, and have also been considered as a target for anti-fungals ^10^. Kinesin-5 family members are found in most eukaryotes, including *Plasmodium sp*., and the family is predicted to have been established in the last eukaryotic common ancestor ^11^. Kinesin-5s from several species form a tetrameric bipolar structure, with two opposing pairs of motor domains that can organise MT arrays such as those found in spindles ^12–14^. Several classes of *Homo sapiens* kinesin-5 (*Hs*K5) inhibitors have been characterised that block motor ATPase activity and bind to allosteric sites in the motor domain. The best studied of these allosteric binding pockets is defined by kinesin-5-specific sequences in a key structural region of the motor domain, loop5 ^15^. Loop5 is critical for the correct operation of human kinesin-5, since its deletion or mutation disrupts the motor’s mechanochemical cycle ^16–18^. Furthermore, *Drosophila* kinesin-5 is resistant to the *Hs*K5 inhibitor STLC, but can be sensitised by replacement of loop5 with the cognate *Hs*K5 sequence ^19^. The loop5-defined allosteric site thus has proven promise in mediating selective inhibition of kinesin-5 family members from different species.

To investigate the idea that kinesin-5 from *Plasmodium falciparum* (*Pf*K5) *-* the deadliest form of human malaria - could be a selective anti-malarial therapeutic target, we characterised the biochemical properties and MT-bound structure of the *Pf*K5 motor domain. We show that the *Pf*K5 motor domain is an ATPase with MT plus-end directed motility, as demonstrated in MT gliding experiments. MT-bound structures of the *Pf*K5 motor domain, determined using cryo-electron microscopy (cryo-EM), reveal classical kinesin mechanochemistry despite the significant sequence divergence of this motor. This includes ATP-induced neck-linker docking to the motor domain, which is consistent with *Pf*K5 motor domain plus-ended directed motility. Finally, we show that a large insertion in loop5 of the *Pf*K5 motor domain creates a dramatically different chemical environment in the well characterised loop5 drug-binding site, revealing the possibility for selective inhibition of *Pf*K5.

## RESULTS

### *Pf*K5ΔL6-MD is a slow ATPase

To characterise *Pf*K5 mechanochemistry, we first wanted to measure its MT-stimulated ATPase activity. The *Pf*K5 motor domain (*Pf*K5MD, amino acids 1-493) contains a 105 amino acid asparagine and lysine rich insertion in loop6 that is characteristic of malaria proteins ^20^ (Fig 1A, left), but which is poorly conserved (16-30% sequence identity) among *Plasmodium* kinesin-5s. To enable our study, we engineered loop6 out of our construct, an approach previously taken by another group ^21^. We refer to this construct as *Pf*K5ΔL6-MD, and it was purified to 99 % purity (Fig 1A, right).

**Figure 1.**
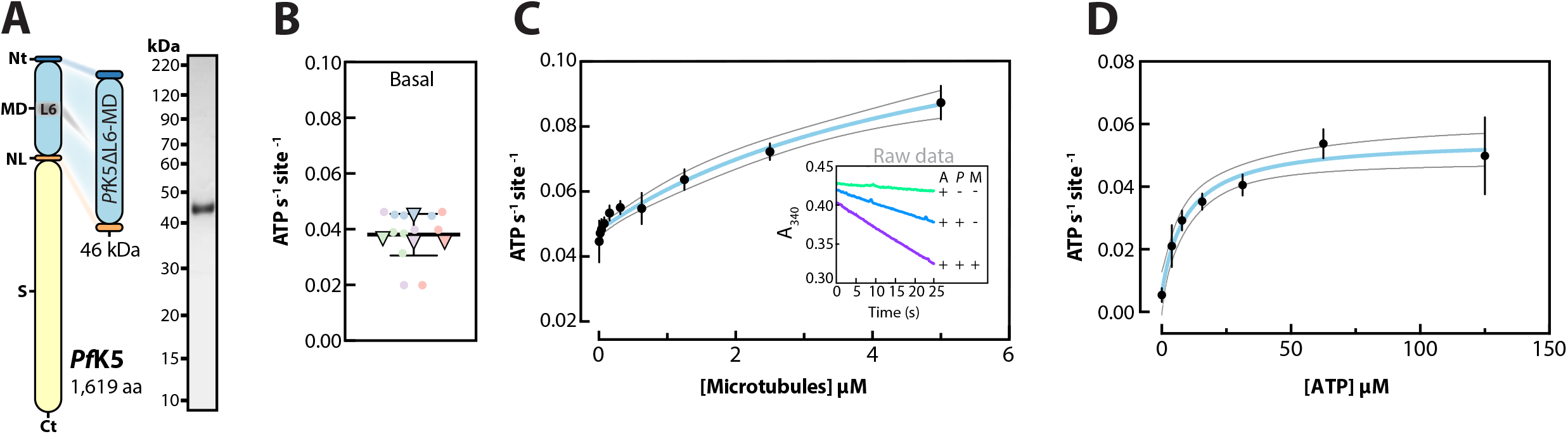
*Pf*K5ΔL6-MD is a slow MT-stimulated ATPase. (A) Left, Domains of full-length *Pf*K5 and *Pf*K5ΔL6-MD, displaying the N-terminus (Nt), motor domain (MD), neck linker (NL), stalk (S), C-terminus (Ct) aa = amino acids; right, Coomassie-stained SDS-PAGE of *Pf*K5ΔL6-MD after purification. (B) *Pf*K5ΔL6-MD ATPase rate in the absence of MTs. Technical replicates = 12 (circles), experimental replicates = 4 (triangles), biological replicates (i.e. number of different protein purifications used in the experiments) = 2. The mean of experimental replicates (0.039 ATP s^-1^) and 95 % confidence interval are plotted. (C) *Pf*K5ΔL6-MD MT stimulated ATPase activity. The mean and standard deviation of 3 experimental replicates (no technical replicates), are plotted. Biological replicates = 2. The fit is plotted as a blue line, with corresponding 95 % confidence interval plotted as black lines. Inset displays an example of raw data (A = ATP, *P* = *Pf*K5ΔL6-MD, and M = MTs). (D) *Pf*K5ΔL6-MD MT stimulated ATPase activity as in (C), except with ATP as the substrate variable, and using a constant of 1 μM MTs.

In the absence of MTs, *Pf*K5ΔL6-MD exhibited a low ATP hydrolysis rate (Fig 1B), but addition of MTs stimulated *Pf*K5ΔL6-MD ATPase activity (Fig 1C). Changes in pH or ionic strength showed no or minimal impact respectively on *Pf*K5ΔL6-MD ATPase rate (S1A,B Fig). From these data, the motor K_cat_ and K_m_ of MTs (K_MT_) for *Pf*K5ΔL6-MD was calculated. The K_m_ of ATP (K_ATP_) was also determined (Fig 1D). *Pf*K5ΔL6-MD has a MT-stimulated ATP hydrolysis rate of 0.13 ATP s^-1^, which is slow compared to kinesin-5 from *S. pombe* ^22^, *S. cerevisiae* ^23^, and *H. sapiens* ^24^ (with rates of 1.2, 0.5, and 2.9 ATP s^-1^ respectively). However, *Pf*K5ΔL6-MD has similar K_MT_ (5.4 μM) and K_ATP_ (9.5 μM) values compared to previously characterised kinesin-5s ^23,24^. Thus, despite substantial sequence divergence, and although it has the lowest ATPase rate observed to date for the family, *Pf*K5ΔL6-MD exhibits overall similar ATPase properties compared to other kinesin-5s.

### *Pf*K5ΔL6-MD generates slow MT gliding

To determine the motile properties of *Pf*K5ΔL6-MD, we used a MT gliding assay. *Pf*K5ΔL6-MD was expressed with a C-terminal SNAP-tag (*Pf*K5ΔL6-MD-SNAP), purified to 92 % purity (Fig 2A), covalently labelled with biotin, and attached to a neutravidin coated surface. The velocity of fluorescently-labelled MTs driven by *Pf*K5ΔL6-MD-SNAP activity was measured. *Pf*K5ΔL6-MD-driven MTs moved at an average velocity of 5.4 nm/s (95 % confidence interval = 5-5.9) (Fig 2A, S1C Fig). This is slow compared to 23-92 nm/s reported for *Hs*K5-MD ^22,25,26^. However, this slow MT gliding corresponds with the slow rate of ATP hydrolysis observed in the ATPase assay. Inclusion of polarity-marked MTs in the assay further showed that *Pf*K5ΔL6-MD is a plus-end directed motor (Fig 2B).

**Figure 2.**
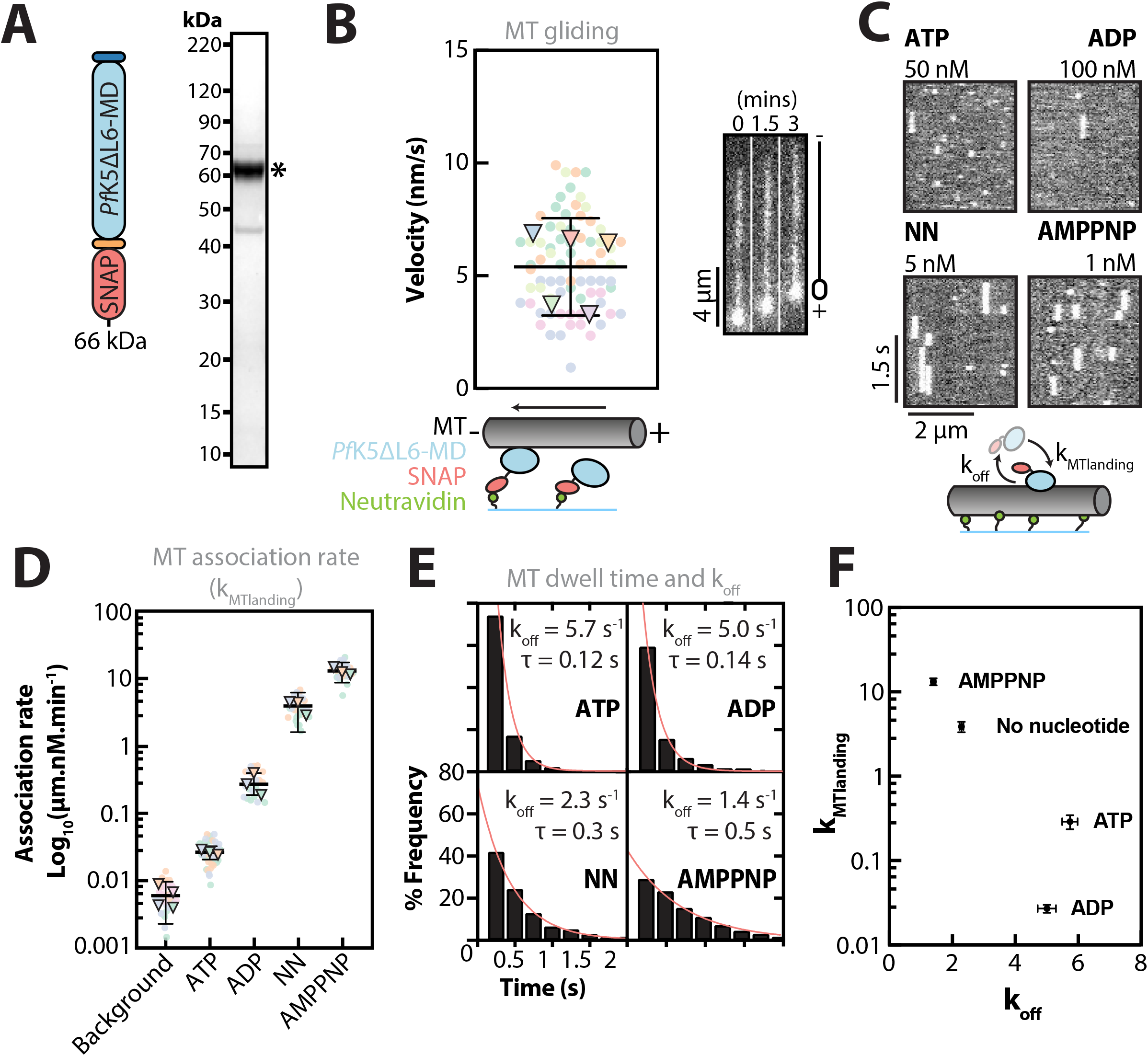
*Pf*K5ΔL6-MD drives plus-end directed MT gliding and its MT interactions are nucleotide modulated. (A) Schematic of the *Pf*K5ΔL6-MD-SNAP fusion protein used for TIRFM experiments, and SDS-PAGE of *Pf*K5ΔL6-MD-SNAP after purification. ^*^ indicates the band for *Pf*K5ΔL6-MD-SNAP. (B) *Pf*K5ΔL6-MD-SNAP driven MT gliding velocity. Technical replicates = 75 (circles), experimental replicates = 5 (triangles), biological replicates = 3. The mean of experimental replicates (5.4 nm/s) and 95 % confidence interval are plotted. Greyscale images on the right are snap-shots of a single bright plus-end labelled MT over time. (C) Example kymographs from *Pf*K5ΔL6-MD-SNAP single molecule MT binding experiments, with each vertical white streak corresponding to a single *Pf*K5ΔL6-MD-SNAP binding event. Concentrations refer to the amount of *Pf*K5ΔL6-MD-SNAP required to see single molecule binding for each nucleotide. (D) *Pf*K5ΔL6-MD-SNAP MT association rates in different nucleotide states (NN = no nucleotide). For each nucleotide condition, technical replicates (circles) and experimental replicates (triangles), are plotted, in addition to the mean and 95 % confidence interval of experimental replicates (Table 1). (E) Frequency distribution of *Pf*K5ΔL6-MD-SNAP MT dwell times, in different nucleotide states. Number of experimental replicates = 3, however frequency distributions are calculated from pooled experimental data (number of technical replicates is shown in Table 1). Number of biological replicates = 2. The fit for one-phase exponential decay models is shown, with corresponding decay constant (k_off_), and the mean dwell time (τ) of *Pf*K5ΔL6-MD-SNAP binding events, which is equal to the half-life of the model. (F) Mean MT association (k_MTlanding_) as a function of MT dissociation (k_off_), plotted with 95 % confidence intervals.

### *Pf*K5ΔL6-MD single molecule interactions with MTs

To better understand the slow ATPase and MT gliding activity we observed for *Pf*K5ΔL6-MD, we analysed the interactions of single molecules of fluorescently labelled *Pf*K5ΔL6-MD-SNAP with MTs. We made single molecule measurements in different nucleotide conditions, to investigate how MT affinity changes with nucleotide state (Fig 2C, S1D Fig, and Table 1), using the non-hydrolysable ATP analogue AMPPNP to mimic the ATP bound state. From these data, we calculated the MT association rate, or k_MTlanding_ (Fig 2D), and MT dissociation rate, or k_off_ of *Pf*K5ΔL6-MD (Fig 2E). This demonstrated that in saturating ADP conditions, *Pf*K5ΔL6-MD-SNAP had a comparatively low kMTlanding and a high k_off_, indicating that *Pf*K5ΔL6-MD-SNAP has low MT affinity when bound to ADP (Fig 2F). In the absence of nucleotide or in saturating AMPPNP conditions, *Pf*K5ΔL6-MD-SNAP had a comparatively high kMTlanding and a low k_off_, showing that when no nucleotide is present, or in its ATP-bound state, *Pf*K5ΔL6-MD-SNAP has high MT affinity.

**Table 1.** Table 1: *Pf*K5ΔL6-MD-SNAP microtubule landing rate and mean microtubule dwell time. In saturating ATP conditions, the average MT dwell time of *Pf*K5ΔL6-MD-SNAP is 0.12 s (Fig 2E) – similar to the ATPase rate of *Pf*K5ΔL6-MD (0.13 ATP s^-1^). This indicates that *Pf*K5ΔL6-MD ATPase activity is rate limited by one of the MT-bound stages of the ATPase cycle. Given that a similar dwell time to the ATPase rate was also observed in ADP saturating conditions (0.14 s), it is therefore possible that *Pf*K5ΔL6-MD is rate limited by MT release when bound to ADP. Taken together, single molecule measurements of *Pf*K5ΔL6-MD-SNAP MT binding support observations of slow ATPase and MT gliding activity.

| Nucleotide State | Microtubule Landing Rate |  |  | Mean Microtubule Dwell Time |  |  |  |
| --- | --- | --- | --- | --- | --- | --- | --- |
| | Number of MTs | $K_{MT\text{landing}}/\text{Rate}$<br>($\mu\text{m.nM.min}^{-1}$) | Std. Dev. | Number of events | $k_{\text{off}}$<br>($\text{s}^{-1}$) | Mean Dwell Time (s) | $R^2$ |
| Background | 34 | 0.006 | 0.0027 |  |  |  |  |
| ATP | 22 | 0.29 | 0.12 | 1065 | 5.7 | 0.12 | 0.99 |
| ADP | 46 | 0.027 | 0.009 | 1084 | 5.0 | 0.14 | 0.99 |
| No nucleotide | 28 | 3.86 | 1.4 | 1799 | 2.3 | 0.30 | 0.99 |
| AMPPNP | 29 | 13.28 | 3.4 | 1717 | 1.4 | 0.50 | 0.99 |

### MT-bound *Pf*K5ΔL6-MD structure determination using cryo-EM

To gain molecular insight into the behaviour of *Pf*K5ΔL6-MD, its interaction with MTs and its sensitivity to nucleotide binding, we visualised MT-bound *Pf*K5ΔL6-MD in different nucleotide states using cryo-EM (Table 2). We calculated 3D reconstructions of *Pf*K5ΔL6-MD bound to MTs in the absence of nucleotide and in an ATP-like state, using AMPPNP. To do this we used MiRP (S2A Fig), our previously developed pipeline for image processing of MTs with RELION ^27,28^. As part of the current study, we have updated MiRP to operate with RELION v3.1, and improved usability of the procedure such that it can be run from the RELION GUI, amongst other updates (see Methods and https://github.com/moores-lab/MiRPv2).

**Table 2.** Table 2: Cryo-EM data collection, 3D Image Processing and model refinement statistics. *Pf*K5ΔL6-MD binds MTs every 8 nm (one *Pf*K5ΔL6-MD per αβ-tubulin) on the ridge of MT protofilaments (Fig 3A), a binding site shared by all other kinesins characterised to date ^29,30^. MiRP initially produced reconstructions for the no nucleotide (Fig 3B) and AMPPNP states (Fig 3C) at overall resolutions of 4.8 Å and 4.5 Å respectively. The resolution in the MT region of both reconstructions is approximately 4.5-5.5 Å - in contrast, the resolution of *Pf*K5ΔL6-MD exhibits a marked falloff as a function of distance from the MT surface (S2B Fig). This is typical of kinesin-MT reconstructions ^31,32^, and is attributable to a number of factors, including incomplete occupancy of the MT lattice by the motor that is not immediately apparent in the micrographs, as well as flexibility in the *Pf*K5ΔL6-MD protein itself. We therefore used symmetry expansion and 3D classification focused on *Pf*K5ΔL6-MD from one *Pf*K5ΔL6-MD:αβ-tubulin dimer asymmetric unit to select data with the best *Pf*K5ΔL6-MD occupancy (S2A Fig). The resulting 3D structures (no nucleotide – Fig 3D, AMPPNP – Fig 3E) were substantially improved, as shown by reduced resolution decay in the motor domain density (S2B,C Fig). The average resolution of these reconstructions was 4.4 and 4 Å for the no nucleotide and AMPPNP states respectively, with *Pf*K5ΔL6-MD having a resolution range of 4.5-7 Å.

|  | *Pfk5ΔL6*-MD nucleotide state |  |
| --- | --- | --- |
|  | No Nucleotide | AMPPNP |
| **Data Collection and 3D Image Processing** |  |  |
| Magnification | 160,000 | 160,000 |
| Voltage (kV) | 300 | 300 |
| Electron dose ($\text{e}^-/\text{\AA}^2$) | 58 | 58 |
| Pixel size ($\text{\AA}$) | 1.39 | 1.39 |
| Number of Images | 1320 | 335 |
| Data Collection Strategy | manual and automated | manual |
| Starting number of particles | 48,165 | 37,843 |
| Final particle number (after particle symmetry expansion) | 166,375 | 73,684 |
| Box size (pixels) | 432 | 432 |
| C1 reconstruction resolution ($\text{\AA}$) | 4.5 | 4.8 |
| Asymmetric unit refine resolution (Å) | 4 | 4.4 |
| **Model Refinement** |  |  |
| Global Cross-correlation: |  |  |
| Homology model | 0.88 | 0.86 |
| Final model | 0.89 | 0.89 |
| MolProbity: |  |  |
| Homology model | 3.46 | 3.46 |
| Final model | 1.13 | 0.98 |
| QMEAN |  |  |
| Homology model | -2.21 | -2.21 |
| Final model | -0.36 | 0.16 |

**Figure 3.**
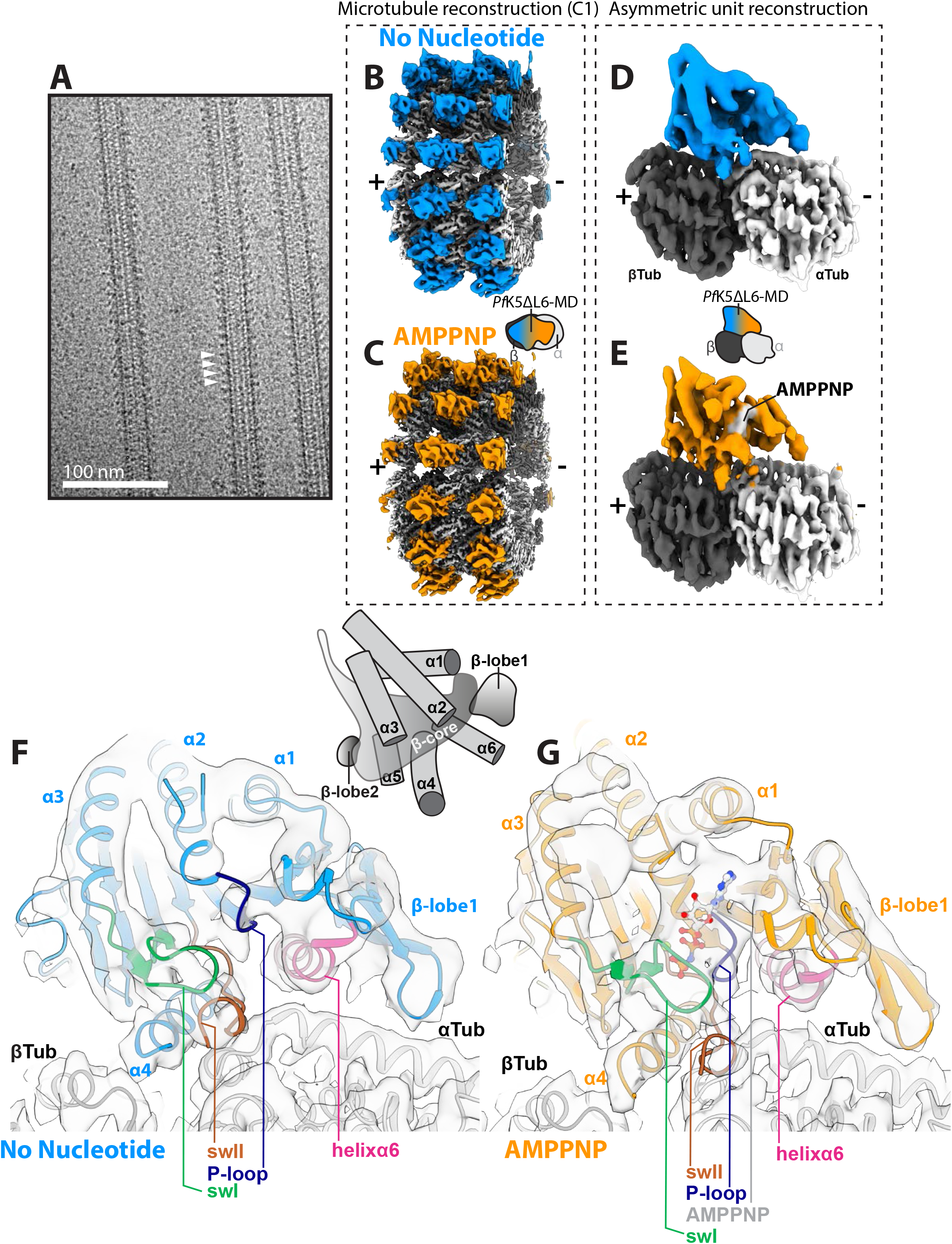
Cryo-EM 3D reconstruction of *Pf*K5ΔL6-MD MT complexes. (A) Example micrograph of the *Pf*K5ΔL6-MD bound MTs with 5 mM AMPPNP. White arrows indicate *Pf*K5ΔL6-MD decoration present every 8 nm (1 αβ-tubulin dimer). (B-E) 3D reconstructions have been locally low-pass filtered according to local resolution. (B) The unsymmetrised (C1) reconstruction of the *Pf*K5ΔL6-MD no nucleotide state bound to MTs, depicting the central portion of the MT reconstruction. (C) as in (B), for the AMPPNP state. (D) Reconstruction of the *Pf*K5ΔL6-MD no nucleotide state bound to αβ-tubulin after asymmetric unit refinement. (E) As in (D), for the AMPPNP state. (F) Ribbon depiction of the no nucleotide state model in corresponding cryo-EM density. (G) As in (F), for the AMPPNP state. In (F) and (G), key components of the *Pf*K5ΔL6-MD are labelled and colour-coded as indicated.

Alignment and superposition of our reconstructions directly reveals nucleotide-dependent differences in these structures (S3A Fig). However, to facilitate interpretation of the differences, we also calculated a *Pf*K5ΔL6-MD homology model and performed flexible fitting of this model within the cryo-EM density, in which individual secondary structure elements were well resolved (S3B Fig). Local cross-correlation scoring showed overall improvement of model fit to density as a result of flexible fitting (S3C Fig), and produced molecular models of each of the *Pf*K5ΔL6-MD:αβ-tubulin dimer complexes (Table 2). These provide a detailed picture of how *Pf*K5ΔL6-MD interacts with both α- and β-tubulin. They also show that this divergent parasite motor has a canonical kinesin fold: it is built from a central β-sheet, sandwiched between three α-helices on each side, and flanked by a small β-sheet (β-lobe1), and a β-hairpin (β-lobe2) (Fig 3F,G). Using these models, we analysed conformational differences between the no nucleotide and AMPPNP states.

### AMPPNP binding causes *Pf*K5ΔL6-MD nucleotide binding site closure

The *Pf*K5ΔL6-MD nucleotide binding site (NBS) is located away from the MT surface, and despite the overall low sequence conservation of *Pf*K5ΔL6-MD compared to *Hs*K5 (S4 Fig), is composed of three loops containing conserved sequence motifs. These are the P-loop - which interacts with the α- and β-phosphates of bound nucleotide - loop9 and loop11, which contain the switch-I and switch-II motifs respectively ^33^.

In the absence of bound nucleotide, density for all three NBS loops is visible in *Pf*K5ΔL6-MD, although density for the P-loop in the no nucleotide state is poorly defined (Fig 4A). In addition, density corresponding to the C-terminal end of loop11, which is approximately 19 Å from the NBS, and contains a two residue *Plasmodium-*conserved insertion, is not visible. In the no nucleotide state, density corresponding to loop9 and the P-loop are well separated, while density is observed connecting loop9 and 11 (Fig 4A). These configurations create an ‘open’ nucleotide binding site primed for ATP binding formed by a cavity between these three loops. In contrast, in the AMPPNP state, there is clear density corresponding to the bound nucleotide in the *Pf*K5ΔL6-MD NBS (Fig 4A, S4 Fig). In addition, the nucleotide is surrounded by loop9 and 11, which have closed around the bound nucleotide to form an NBS that supports ATP hydrolysis (Fig 4A) ^34^.

**Figure 4.**
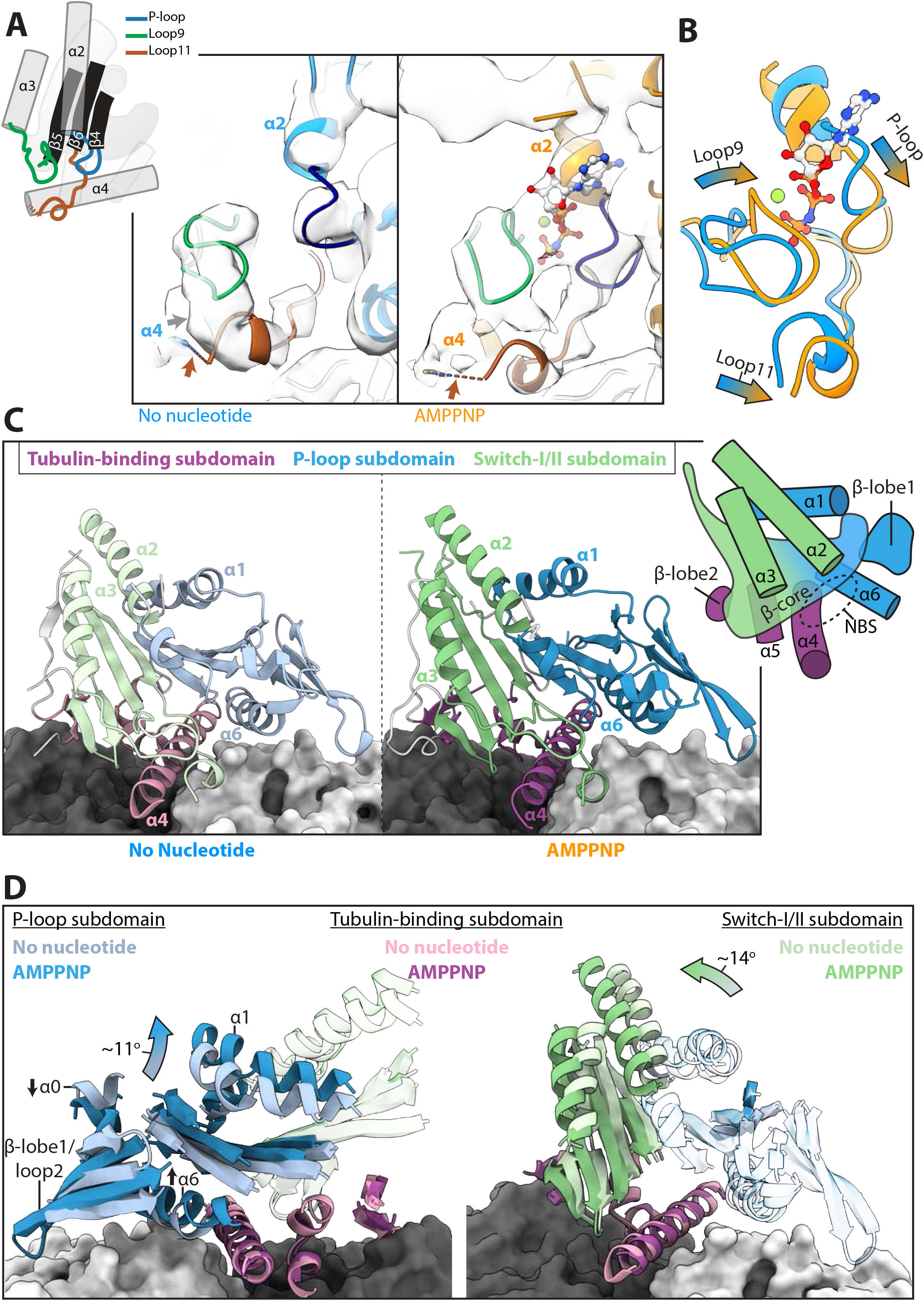
AMPNPP binding causes *Pf*K5ΔL6-MD subdomain rearrangement and switch loop closure. (A) Rearrangements of *Pf*K5ΔL6-MD switch loops upon AMPPNP binding, with a schematic showing connectivity of the switch loops to motor domain secondary structure elements on the left. Models and corresponding density are displayed on the right, with nucleotide binding loops coloured according to the schematic. The brown arrow indicates where there is missing density for loop11, and the grey arrow indicates connecting density between loop9 and 11. (B) Comparison of *Pf*K5ΔL6-MD no nucleotide and AMPPNP nucleotide binding loops, demonstrating AMPPNP induced conformational changes. (C) The *Pf*K5ΔL6-MD no nucleotide state model, coloured according to kinesin subdomain, with α and β-tubulin depicted in light and dark grey surface rendering respectively. (D) *Pf*K5ΔL6-MD subdomain rearrangement, showing no overall movement in the tubulin-binding subdomain, with rotations in the P-loop subdomain shown on the left, and the switch-I/II subdomain on the right.

Superimposing the *Pf*K5ΔL6-MD no nucleotide and AMPPNP state models by alignment on αβ-tubulin allows visualisation of the structural response of the *Pf*K5ΔL6-MD NBS to AMPPNP binding (Fig 4B). Even while the resolution is not sufficient to determine the exact conformation of each of these loops, it is very clear that in the presence of AMPPNP, all three NBS loops move, with loop9 and the P-loop coming closer together, thereby burying the nucleotide. In summary, AMPPNP binding to *Pf*K5ΔL6-MD causes a conformational rearrangement that forms a closed, catalytically competent NBS.

### AMPPNP binding causes *Pf*K5ΔL6-MD subdomain rearrangement

What are the consequences of these NBS rearrangements on the structure of *Pf*K5ΔL6-MD? The structure of *Pf*K5ΔL6-MD can be subdivided into three distinct subdomains ^35^, which are predicted to move with respect to each other during the motor’s MT-based ATPase cycle (Fig 4C). The tubulin-binding subdomain (Fig 4C, purple hues) consists of MT binding elements in which helixα4 binds a shallow cavity at the intra-tubulin dimer interface. In addition, helixα5 and β-lobe2 contact β-tubulin. The remaining two subdomains, the P-loop subdomain (Fig 4C, blue hues) and switch-I/II subdomain (Fig 4C, green hues), contain approximately half of the central β-sheet each, along with adjacent secondary structure elements. The NBS is located at the junction of these subdomains (Fig 4C).

We measured the relative rotation of each helix (S3 Table) in the transition from no nucleotide to AMPPNP states. This reveals the rearrangement of the P-loop and switch-I/II subdomains around a static MT binding domain (Fig 4D). The P-loop subdomain pivots such that helixα0 moves towards the MT surface, while helixα6 and the majority of the subdomain moves away from the MT surface. The switch-I/II subdomain rotates such that its constituent secondary structure elements move towards β-tubulin.

### AMPPNP binding causes *Pf*K5ΔL6-MD neck linker docking

What are the consequences for these subdomain rearrangements for the functional output of *Pf*K5ΔL6-MD? Approximately 18 Å away from the MT surface, and 27 Å away from the NBS, is the neck linker. The neck linker is a C-terminal peptide extending from helixα6, which links the motor domain to the kinesin stalk, and which is relatively conserved in the kinesin superfamily (S3C Fig). In the *Pf*K5ΔL6-MD no nucleotide reconstruction, no density is observed extending from helixα6, indicating that the neck linker is disordered in this nucleotide state (Fig 5A). In the AMPPNP state however, there is clear density for the neck linker at the C-terminus of helixα6 that extends along the motor domain in the direction of the MT plus-end (Fig 5B). In addition, density corresponding to the *Pf*K5ΔL6-MD N-terminus is also visualised in the AMPPNP state, consistent with formation of backbone interactions between this region and the neck linker, to form short β-strands known as the cover neck bundle (CNB) ^36^ (Fig 5B, black arrow; S6A,B Fig). No such CNB density is observed in the no nucleotide reconstruction (S6B Fig). Neck linker docking is enabled by the above-described rotation of the P-loop subdomain, which moves the N-terminus and the central β-sheet away from helixα5/loop13 in the static MT binding subdomain (Fig 4D, Fig 5) ^37,38^. This creates a cavity between loop13 and the N-terminus, known as the docking cleft, which enables neck linker docking. Thus, *Pf*K5ΔL6-MD subdomain rearrangement causes opening of the docking cleft upon AMPPNP binding, a structural transition that is consistent with the ability of *Pf*K5ΔL6-MD to drive ATP-dependent plus-end directed gliding motility ^35^.

**Figure 5.**
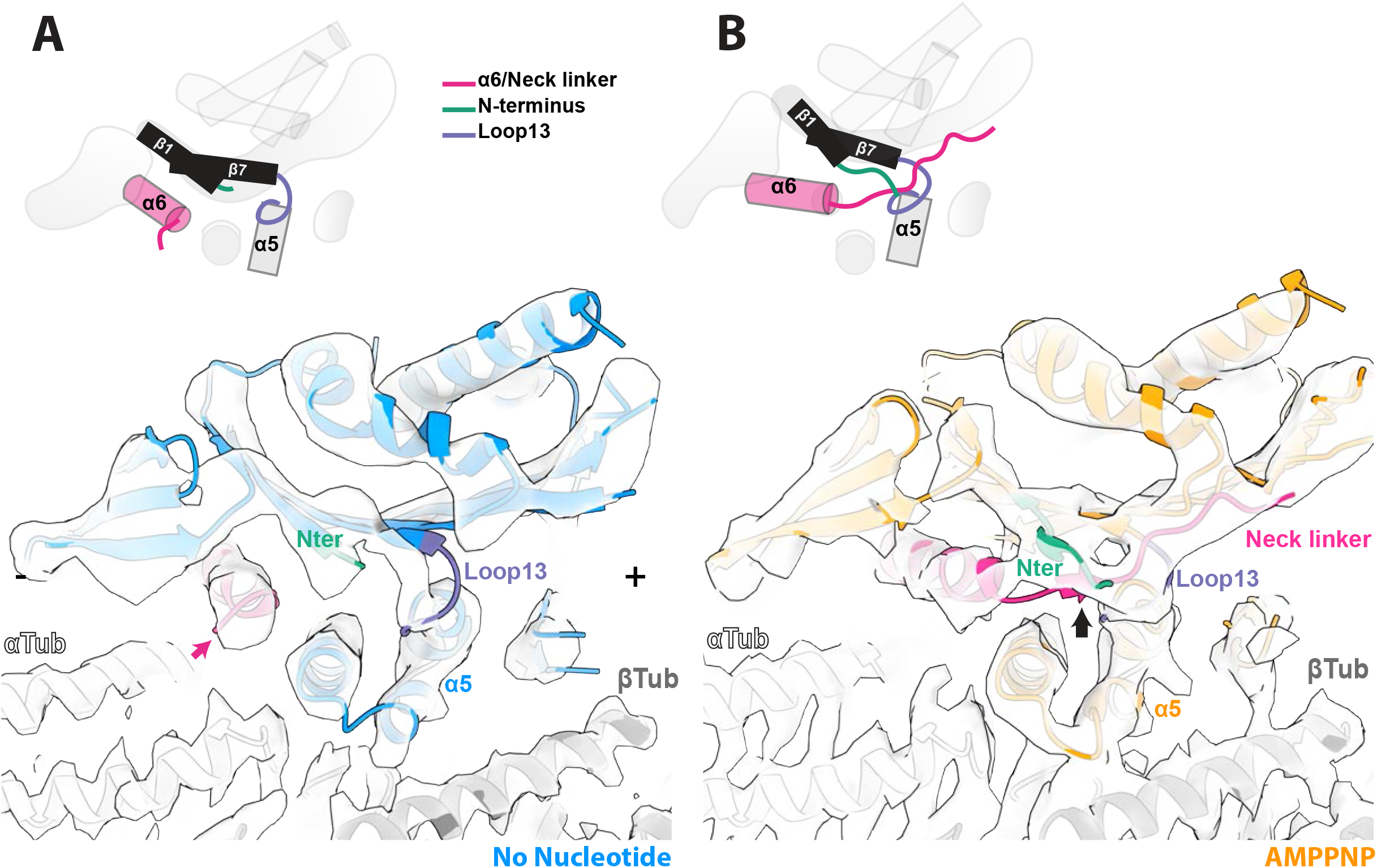
AMPNPP binding causes *Pf*K5ΔL6-MD neck linker docking. (A) No nucleotide *Pf*K5ΔL6-MD model and corresponding cryo-EM density of the neck linker docking site. Helixα6/neck linker, the N-terminus, and loop13 are coloured according to the schematic. Note the lack of density for the neck linker at the terminus of helixα6. +/- symbols denote the MT polarity. Pink arrow indicates the lack of density for an ordered neck linker. (B) as in (A) for the AMPPNP state, showing the neck linker and corresponding density docked along the motor domain.

### MT binding interface

*Pf*K5ΔL6-MD binds to one αβ-tubulin dimer, with helixα4 centred at the intra-dimer interface (Fig 6A). At the *Pf*K5ΔL6-MD binding site, the sequence of the *S. scrofa* αβ-tubulin used in our reconstruction is identical to that of *P. falciparum* αβ-tubulin (S7 Fig), facilitating a more detailed investigation of this interface. To analyse which *Pf*K5ΔL6-MD secondary structure elements interact with αβ-tubulin in the no nucleotide state, we coloured an αβ-tubulin surface representation according to different *Pf*K5ΔL6-MD secondary structure elements; we also coloured *Pf*K5ΔL6-MD according to proximity to α or β-tubulin (Fig 6B). This analysis shows that helixα4 interacts with both α- and β-tubulin, while βlobe1, loop11 and helixα6 interact with helices 4, 5 and 12 of α-tubulin, and loop7, βlobe2 and helixα5 interact with similar secondary structure elements in β-tubulin.

**Figure 6.**
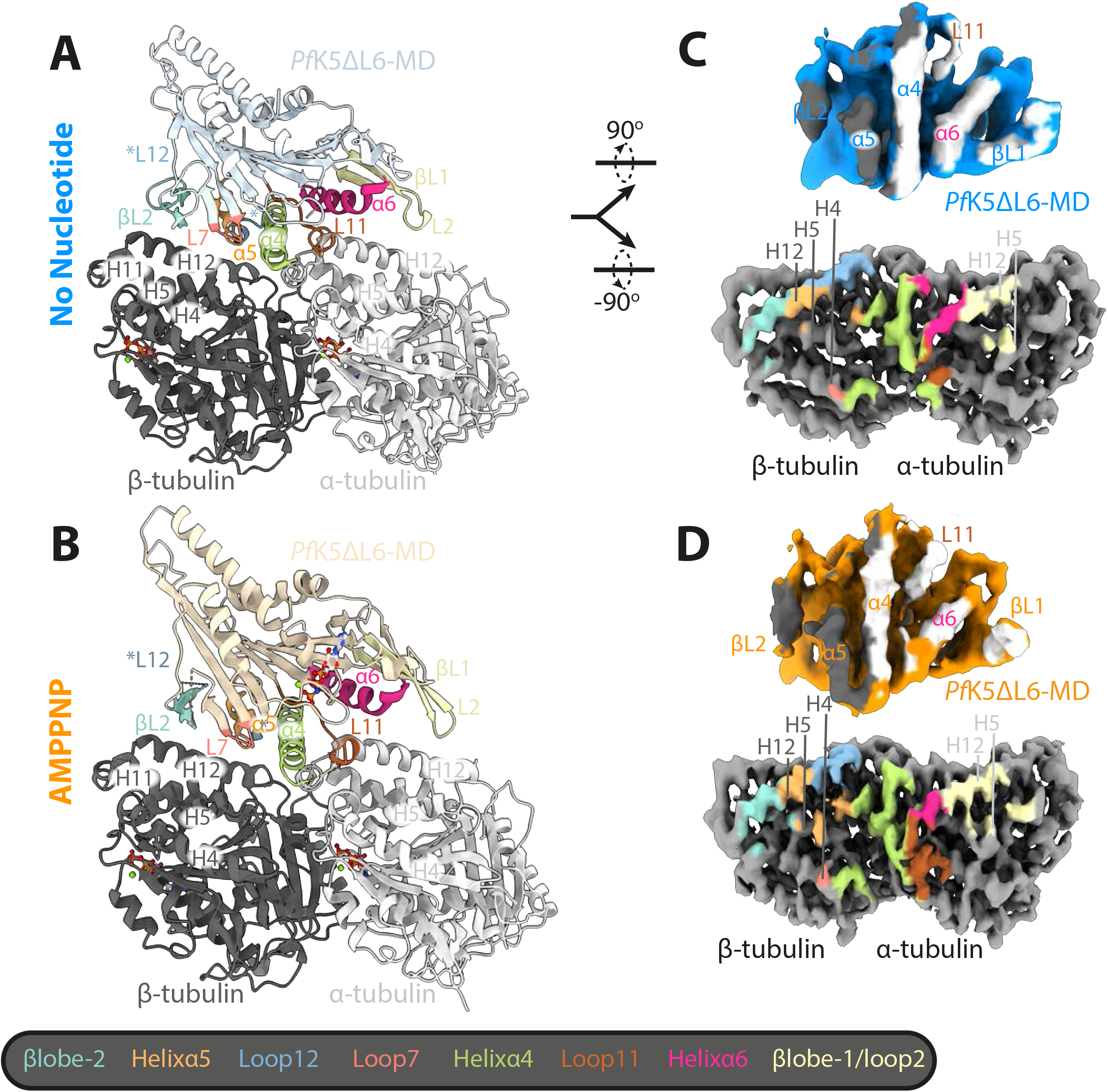
*Pf*K5ΔL6-MDhas an altered MT interface. (A) The *Pf*K5ΔL6-MD no nucleotide state model in ribbon and coloured light blue. with secondary structure elements partaking in the MT interface coloured according to the key below. (B) As for in (A), for the *Pf*K5ΔL6-MD AMMPNP state model, coloured in light orange. (C) Upper image – no nucleotide *Pf*K5ΔL6-MD density, rotated 90^°^ from (A), and coloured according to αβ-tubulin when a particularly area of density is <7 Å away. Lower image - αβ-tubulin density from the no nucleotide state, coloured according to different *Pf*K5ΔL6-MD secondary structure elements, as outlined in the key. (D) As in (C), for the *Pf*K5ΔL6-MD AMPPNP state.

Much of this *Pf*K5ΔL6-MD-MT interface is similar between the no nucleotide and AMPPNP states, (Fig 6C, S4 Table). However, subdomain re-arrangement and NBS closure in the AMPPNP state result in some changes. The largest of these occurs at the α-tubulin interface, where rotation of the P-loop subdomain decreases the interaction of helixα6 with α-tubulin, and positions βlobe1 closer to α-tubulin, increasing its interface area. The interface area of loop11 also increases in the AMPPNP state. Interestingly, βlobe1/loop2 forms an interface area with α-tubulin of 181 and 145 Å^2^ in the no nucleotide and AMPPNP states respectively. Taken together, this shows that the *Pf*K5ΔL6-MD-MT interface is similar to that observed for other kinesin-5s, although the extent to which βlobe1/loop2 interacts with α-tubulin differs between different family members ^10,22,39^.

### *Pf*K5ΔL6-MD loop5 forms a unique putative drug binding site

Loop5 plays an important role in the mechanochemistry of *Hs*K5 ^16,40^, and forms the drug binding pocket of *Hs*K5-specific inhibitors ^19^. It is a solvent exposed loop that creates a break in helixα2, and protrudes from the surface of the motor domain away from the MT. The loop5 sequence is well conserved between *Plasmodium* species (76-91 % sequence identity), and is longer compared to *Hs*K5 (Fig 7A). In the no nucleotide state of *Pf*K5ΔL6-MD, some very poorly defined density corresponding to loop5 can be seen at a low threshold (Fig. 7B), suggesting that this region is largely disordered in this state.

**Figure 7.**
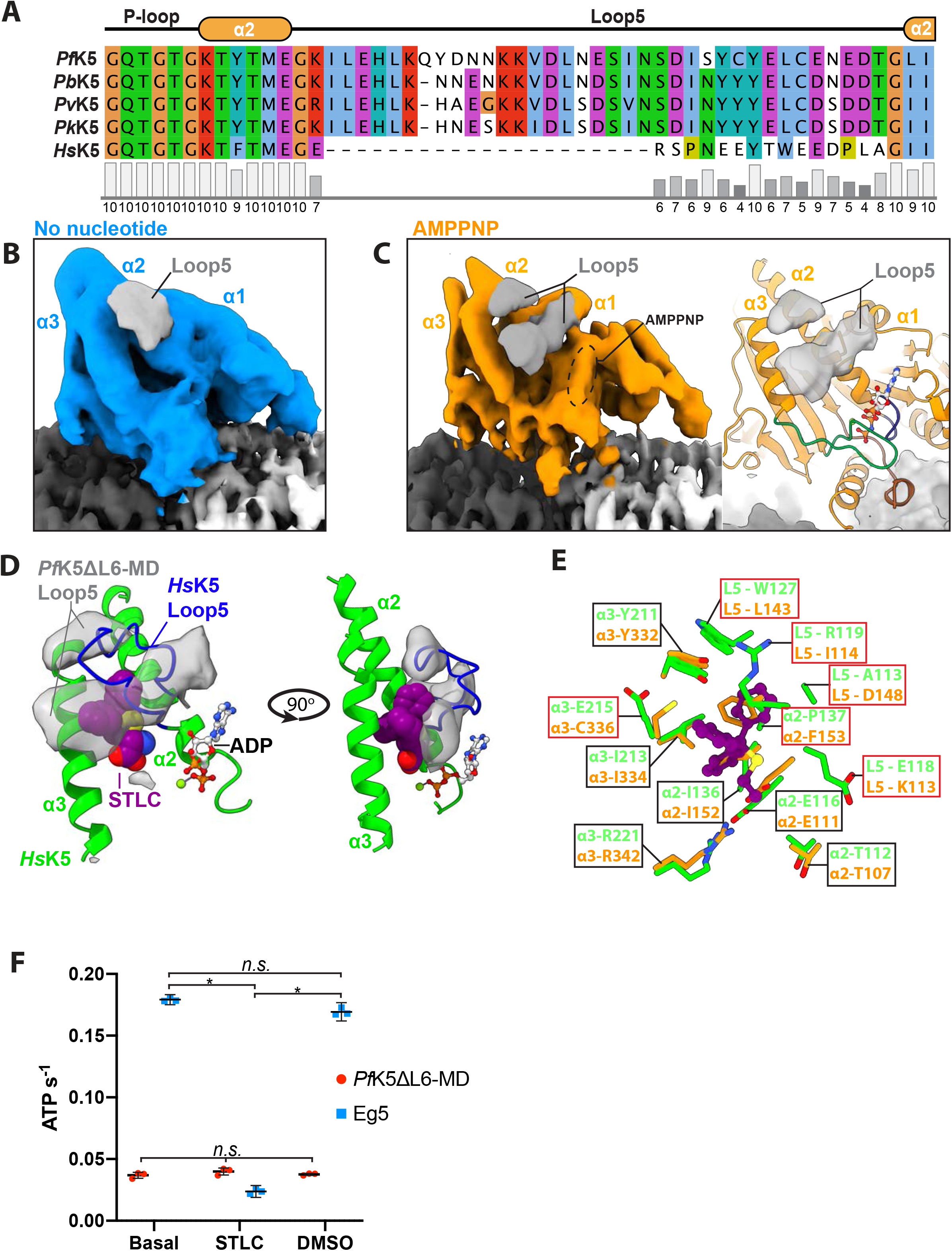
*Pf*K5ΔL6-MD loop5 alters the spatial and chemical environment of the kinesin-5 drug binding site. (A) Primary sequence alignment of loop5 from *Hs*K5 (UniProt ID: P52732), *Pf*K5 (O77382) and various other *Plasmodium* species (*Pv* = *vivax* (A0A564ZV10), *Pk* = *knowlesi* (A0A1Y3DTC2), *Pb* = *berghei* (A0A122I4M3)). Conserved positions are coloured according to the ClustalX scheme, and a conservation score as calculated in Jalview is given below. (B) Loop5 density (grey) in the no nucleotide state, compared to other *Pf*K5ΔL6-MD density (blue). (C) Loop5 density (grey) in the AMPPNP state, compared to other *Pf*K5ΔL6-MD density (left), and the *Pf*K5ΔL6-MD model (right). (D) The STLC bound *Hs*K5 crystal structure in lime (PDB ID: 2WOG ^41^), which was rigid body fitted into the *Pf*K5ΔL6-MD AMPPNP state map. STLC is coloured purple, and *Hs*K5 loop5 is coloured blue. *Pf*K5ΔL6-MD loop5 cryo-EM density is depicted in grey. (E) Conservation of residues partaking in STLC binding between *Pf*K5ΔL6-MD and *Hs*K5. *Hs*K5 residues contacting STLC were found using Chimera, and are shown with equivalent *Pf*K5ΔL6-MD residues. Non-conserved residues are displayed in red boxes. (F) ATPase rates of *Pf*K5ΔL6-MD and *Hs*K5 rate in the absence of MTs, with either no treatment, + 20 μM STLC, or DMSO (with the same % v/v as the % v/v of STLC). Statistical relationships were tested using a one-way ANOVA, followed by a post-hoc Tukey’s multiple comparison test.

In the AMPPNP state, however, clear density corresponding to loop5 can be seen, which forms two distinct regions. This density extends at an angle from helixα2, forming elongated density projecting from the motor domain between helixα1 and helixα3 (Fig 7C). This region is at lower resolution than other parts of the reconstruction, possibly owing to intrinsic flexibility, but also possibly because of the residual resolution gradient towards the outer surface of *Pf*K5ΔL6-MD (S2B Fig). There is no secondary structure-like density in this region of the motor, consistent with sequence-based predictions (S5 Fig) and, therefore, a model for *Pf*K5ΔL6-MD loop5 was not be calculated.

Strikingly, however, the density corresponding to loop5 in the AMPPNP state does not protrude away from the surface of the motor but appears to cover the site between helicesα2 and 3, equivalent to the well described inhibitor binding site in *Hs*K5. Docking of a crystal structure of *Hs*K5 bound to the well-characterised inhibitor STLC in the *Pf*K5ΔL6-MD density reveals the poor match between *Hs*K5 loop5 and the *Pf*K5ΔL6-MD loop5 density (Fig 7D). This also suggests that, although residues outside of loop5 involved in interactions with STLC are largely conserved between *Hs*K5 and *Pf*K5 (Fig 7E), loop5 of *Pf*K5ΔL6-MD might radically alter the spatial and chemical environment of this putative drug binding site. To test this idea, we measured whether the ATPase activity of *Pf*K5ΔL6-MD was susceptible to inhibition by STLC ^42^. Consistent with our structural prediction, while STLC inhibits *Hs*K5 ATPase activity, it does not inhibit *Pf*K5ΔL6-MD (Fig 7F). Thus, despite the conserved aspects of *Pf*K5ΔL6-MD mechanochemistry uncovered by our data, evolutionary divergence between *Pf*K5ΔL6-MD and *Hs*K5 mediates differential inhibition of these kinesin-5 motors.

## DISCUSSION

We have determined the biochemical properties and MT-bound cryo-EM structures of a spindle-associated kinesin-5 motor from the malaria parasite. Despite considerable divergence from the human host kinesin-5 sequence, our *P. falciparum* kinesin-5 *Pf*K5ΔL6-MD construct shares with *Hs*K5 a comparatively slow MT-stimulated ATPase, plus-end directed MT gliding activity and nucleotide-dependent conformational changes that support plus-end directed motility. Significantly, however, our structures revealed a radically different configuration of the well characterised loop5-defined drug binding pocket. Further, we also showed that *Pf*K5ΔL6-MD exhibits no sensitivity to the classical *Hs*K5 loop5 binding drug STLC.

The steady state ATPase activity of *Pf*K5ΔL6-MD is ∼340 times slower than *H. sapiens* kinesin-1 ^38^, 3-25 times slower than other members of the kinesin-5 family ^23,24,39^, and its MT gliding activity is similarly and proportionally slow. Our use of mammalian brain tubulin rather than native *P. falciparum* tubulin might, in principal, contribute to this - however, αβ-tubulin is well conserved between *S. scrofa* and *P. falciparum*, and the two species have identical residues at the kinesin binding site (S7 Fig), suggesting that tubulin source is unlikely to influence *Pf*K5ΔL6-MD activity. In further support of this, experiments comparing ATPase rates of a yeast kinesin-5 motor domain interacting with mammalian and yeast tubulin showed no difference ^39^. A previous study of *Plasmodium falciparum* and *vivax* kinesin-5s also observed slow ATPase rates for these motors ^21^. Our single molecule MT binding data suggest that *Pf*K5ΔL6-MD ATPase activity is rate limited during the MT-bound portion of its ATPase cycle. These findings are reminiscent of the properties of other kinesin-5s and indeed, this may be critical for their function – substitution of the slow motor activity of vertebrate kinesin-5 with faster kinesin-1 was functionally disruptive in the complex context of the spindle ^43^. This suggests that *Pf*K5 – like other kinesin-5s – operates in motor ensembles, where slow-moving teams of *Pf*K5 collaborate to drive MT organisation ^13^.

The malaria kinesin-5 protein we studied was engineered to remove a low-complexity region in loop6, a strategy that had previously been adopted both in characterising malaria kinesin-5 ^21^ and other malaria proteins ^44^. The insertion point of loop6 lies approximately 40 Å from the NBS and, although we cannot exclude that removal of this region influences *Pf*K5ΔL6-MD’s behaviour, our structures clearly demonstrate that the engineered protein adopts a canonical kinesin fold and undergoes a structural response to AMPPNP binding. Loop6 residues are therefore not required for protein folding and fundamental kinesin mechanochemistry. Low-complexity regions like *Pf*K5 loop6 are very common in malaria proteins, and are often found inserted in otherwise well-conserved three-dimensional folds ^20^. While the role of such low-complexity regions in immune evasion is logical for extracellular parasite proteins ^45^, it remains unclear if and how such regions modulate intracellular protein function.

Improvements of our MiRP image analysis procedures allowed us to efficiently handle the incomplete binding of *Pf*K5ΔL6-MD along the MTs in our cryo-EM data (S2A Fig) and to clearly visualise MT-bound *Pf*K5ΔL6-MD at 5-6 Å resolution. Our structures showed that *Pf*K5ΔL6-MD exhibits an open-to-closed conformational change in the NBS, kinesin motor subdomain rearrangements and neck linker docking on ATP analogue binding, typical of a classical plus-end kinesin ^38,46,47^. The lowest resolution region of our *Pf*K5ΔL6-MD is its MT-distal surface, which encompasses the potential drug-binding loop5 region. Because we used GMPCPP-stabilised MTs, we do not think that MT lattice discontinuities – that can occur on paclitaxel-stabilised MTs ^32,48^ – cause this resolution loss. Rather, loop5 of *Pf*K5ΔL6-MD, which is 21 residues longer than in *Hs*K5 and composed mainly of hydrophilic residues, appears to be intrinsically flexible and thus its conformation is more challenging to capture structurally. Density for loop5 is only well defined in the AMPPNP state and not the no nucleotide state, suggesting it is conformationally sensitive to bound nucleotide, as also observed with *Hs*K5 loop5 ^49^. Strikingly, density attributable to loop5 impinges on the pocket corresponding to the well-characterised drug binding site in *Hs*K5 ^50^ and provides a possible explanation for the lack of sensitivity of *Pf*K5ΔL6-MD to inhibition by the small molecule STLC. Given the strong sequence conservation in loop5 between different *Plasmodium* species, this encouraging finding raises the possibility of selective inhibition of parasite motors. Indeed, a small molecule screen identified a compound able to inhibit *Plasmodium* kinesin-5 ATPase activity, but not that of *Hs*K5 ^21^.

*P. berghei* kinesin-5 localises to mitotic and meiotic spindles in blood and mosquito stages of the parasite life cycle ^51^, consistent with a conserved role for this motor in the parasite cell division machinery. Although we know very little else about the function of this motor, we infer from our biochemical and structural data that *Plasmodium* kinesin-5 is likely to play a MT-organising role within parasite spindles. Kinesin-5 is not essential during the blood stages of the *Plasmodium* life cycle ^51,52^. However, knockout of *P. berghei* kinesin-5 substantially reduces the number of sporozoites in oocysts and mosquito salivary glands. This highlights the operational diversity of replication at different parasite life cycle stages in general, and specifically suggests a key role for kinesin-5 in the multiple rounds of mitosis that occur during sporozoite production in the mosquito host ^51^.

There is increasing focus on tackling malaria not only during the symptomatic blood stages of the parasite life cycle but also by perturbing *Plasmodium* transmission between vector and host to facilitate malaria control at the population level ^53^. Intriguingly, despite the reduction of sporozoite numbers in kinesin-5 knockout parasites, the residual sporozoites achieved normal infectivity. Nevertheless, the role of kinesin-5 in this life cycle stage sheds light on parasite transmission vulnerabilities. Particularly given the distinct parasite number threshold that supports onward transmission between vector and host ^54^, combinations of perturbations in sporozoite production could enable transmission control. Moreover, the diverse mechanisms by which small molecules can inhibit *Hs*K5 function have demonstrated that some modes of motor inhibition can be more functionally disruptive than preventing MT binding or than removing motor function completely, for example by trapping it in a tightly bound MT state. In fact, tight MT binding is the proposed mechanism for the anti-fungal small molecules that target *C. albicans* kinesin-5, despite that motor being non-essential ^55^. In the context of these promising findings, our data provide a structural basis for future investigations into parasite-specific kinesin inhibitors.

## MATERIALS AND METHODS

### Protein expression and purification

The *Pf*K5 motor domain (*Pf*K5MD, residues 1-493) was engineered such that 105 amino acids of the asparagine/lysine rich insertion in loop6 (residues 175-269) were removed to facilitate protein expression. The resulting construct, which we refer to as *Pf*K5ΔL6-MD, was cloned in a pET-151D-TOPO vector (Invitrogen) with an N-terminal His_6_-tag and TEV protease cleavage site, and each preparation was expressed in 12 L BL21 Star™DE3 *E. coli* cells (Invitrogen) grown in LB media. Cells were grown at 37^°^ C until they reached an optical density of 0.8-1.0, and were then induced with 0.1 mM IPTG for 3 hours at 26^°^C. Cells were harvested (6,300 g, 15 mins, 4^°^C) and stored at -80^°^C.

Cells were lysed with 3 passages through a C3 homogeniser (Avestin) in 100 mL IMAC W buffer (50 mM Tris-HCl pH 8, 400 mM NaCl, 2 mM MgCl2, 2 mM DTT, 1 mM ATP, 10 mM imidazole pH 8), which was supplemented with 2 x cOmplete™ EDTA-free protease inhibitor tablets (Roche), 15 μg/mL DNase I (Roche), 0.5 mg/mL Lysozyme, 10 % v/v glycerol. The lysate was then clarified by centrifugation (48,000 g, 45 mins, 4^°^C). *Pf*K5ΔL6-MD was purified from the clarified lysate at 4^°^C using an ÄKTA Pure (GE Healthcare) in one day, which reduced loss of protein due to aggregation. First, nickel affinity chromatography was performed where the lysate was loaded onto a 5 mL HisTrap™ Excel column (GE Healthcare), followed by a wash with 20 column volumes (CV) of IMAC W, then reverse elution with 10 CV of IMAC E buffer (same composition as IMAC W, but with 300 mM imidazole). The eluate from this step was concentrated to 10-15 mL with a Vivaspin^®^ concentrator (Sartorius). Concentrated sample was then exchanged into IEX W buffer (same as IMAC W, except containing 80 mM NaCl and no imidazole) using a HiPrep 26/10 desalting column (GE Healthcare). Next, anion exchange chromatography (1 mL HiTrap™ Q HP) was performed, where the flow-through and wash fractions containing *Pf*K5ΔL6-MD were pooled. The His_6_-tag was then cleaved by incubation with 100 μg/mL TEV protease for 2-4 hours at 4^°^C. TEV-protease and any remaining contaminants were removed using nickel affinity chromatography (1 mL HisTrap™ HP), where flow-through and wash fractions containing *Pf*K5ΔL6-MD were pooled. *Pf*K5ΔL6-MD was concentrated to 100-150 μL then, using 0.5 mL Zeba™ 7K MWCO spin columns, was exchanged into T50K20 buffer (50 mM Tris-HCl pH 8, 20 mM KCl, 2 mM MgCl_2_, 2 mM DTT). This purified *Pf*K5ΔL6-MD was snap frozen in liquid nitrogen and stored at -80^°^C. *Pf*K5ΔL6-MD purity was measured using Coomassie-stained SDS-PAGE band intensity measurement in Fiji ^56^.

### Protein labelling

Gibson assembly ^57^ was used to clone in-frame a C-terminal SNAP-tag on *Pf*K5ΔL6-MD (*Pf*K5ΔL6-MD-SNAP) for use in total internal reflection microscopy (TIRFM) experiments. Expression and purification of *Pf*K5ΔL6-MD-SNAP was performed as for *Pf*K5ΔL6-MD, except the anion exchange chromatography step was altered as follows: concentrated and desalted sample was added to a 1 mL HiTrap™ Q HP column, washed with 10 CV IEX W buffer, and eluted with a 20 CV gradient elution to 500 mM NaCl. Eluted fractions containing *Pf*K5ΔL6-MD-SNAP were pooled. *Pf*K5ΔL6-MD-SNAP was biotinylated or fluorescently labelled by overnight incubation at 4^°^C with SNAP-Biotin^®^ or SNAP-Surface^®^ Alex Fluor^®^ 647 (New England BioLabs) with at least a 3:1 molar excess of these labels to *Pf*K5ΔL6-MD-SNAP. Free SNAP-ligand was removed by 2 repeats of buffer exchange in T50K20 buffer (0.5 mL Zeba™ 7K MWCO spin columns).

### MT Preparation

Purified and lyophilised unlabelled, X-rhodamine labelled, or biotin labelled *Sus scrofa* brain tubulin - except X-rhodamine which was from *Bos taurus* -(catalogue numbers T240C, TL620M T333P, respectively, all >99% pure, Cytoskeleton Inc.) was reconstituted to 10 mg/mL in BRB80 (80 mM PIPES pH 6.8, 2 mM MgCl_2_ 1 mM EGTA pH 6.8), centrifuged at 611453 g for 10 mins at 4^°^C, and the supernatant snap frozen in liquid nitrogen and stored at -80^°^C. Double cycle GMPCPP polymerisation was performed as follows: reconstituted tubulin was supplemented with 1 mM GMPCPP (Jena Biosciences) and incubated for 5 mins on ice. Tubulin was then polymerised at 4-5 mg/mL for 20 mins at 37^°^C. MTs were then pelleted at 611453 g for 10 mins at 23^°^C, washed twice, then resuspended, both with BRB80. MTs were then depolymerised on ice for 15 mins, and a second round of polymerisation performed as above. MTs for the ATPase assay were pelleted on a 50 % v/v sucrose/BRB80 cushion after the second polymerisation, to aid separation of MTs from unpolymerised tubulin.

For labelled MTs, X-rhodamine and unlabelled tubulin were mixed in a 1:9 ratio, or X-rhodamine, biotinylated, and unlabelled tubulin were mixed in a 1:1:8 ratio and polymerised at approximately 4 mg/mL total tubulin with 1 mM GTP at 37^°^C for 20 mins. 40 μM Paclitaxel (Merck) dissolved in DMSO was then added, followed by a further incubation at 37^°^C for 15 mins, and then incubation at room temperature for at least one day before use. To prepare polarity marked MTs ^58^, long, dimly labelled MTs (1:9 ratio of X-rhodamine to unlabelled tubulin) were polymerised at 2 mg/mL total tubulin for 2 hours with 1 mM GMPCPP. To prepare NEM-tubulin, by which minus-end MT growth is blocked, 8 mg/mL unlabelled tubulin was incubated with 1 mM N-ethyl maleimide (Sigma) on ice for 10 mins, then with 100 mM ß-mercaptoethanol (Sigma) for 10 mins. To polymerise the bright plus end MT cap, NEM-tubulin was mixed 1:1 with X-rhodamine tubulin and incubated at 37 ^°^C for 15 mins. Finally, long, dim MTs were pelleted (15 mins, 17,000 g, room temp.), the pellet resuspended with bright MT caps, incubated 37^°^C for 15 mins, then 40 μM paclitaxel added.

### Steady-state ATPase activity

An NADH-coupled ATPase assay ^59^ containing 1.5 mM Phosphoenolpyruvate (Sigma), 9-15 U/mL pyruvate dehydrogenase (Sigma), 13.5-21 U/mL lactate dehydrogenase (Sigma) and 0.25 mM NADH (Roche) was used to measure *Pf*K5ΔL6-MD ATPase rates. Reactions were performed in T50K40 buffer (50 mM Tris-HCl pH 8, 400 mM KCl, 2 mM MgCl_2_, 2 mM DTT) with 100-200 nM *Pf*K5ΔL6-MD. A_340_ readings were taken on a SpectraMax 384 (Molecular Devices) from 100 μL reactions at 26^°^C, with automatic path length correction to 1 cm, taking readings every 10 s for 30 mins. Background readings containing no *Pf*K5ΔL6-MD were subtracted from all readings. ATP hydrolysis per second per site was derived with the Beer-Lambert equation, and K_cat_ calculated using the Michaelis-Menten equation with the term K_0_ (rate at substrate concentration of 0).

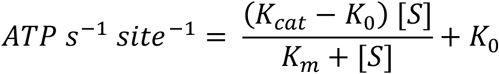

### MT gliding and single molecule TIRFM assays

TIRFM assays with fluorescently labelled protein were performed in a flow-chamber, created by adhering biotin-PEG coverslips (MicroSurfaces Inc.) to glass slides with double-sided tape. To prepare flow-chambers for the MT gliding assay, the following treatments were made. (1) 0.75 % Pluronic acid + 5 mg/mL casein was added for 5 min, and the chamber was washed with T50K20 buffer + 20 μM paclitaxel (T50T). (2) 0.5 mg/mL neutravidin was added for 2 mins, then the chamber was washed twice with T50T + 1 mg/mL casein (T50TC). (3) 1-2.5 μM *Pf*K5ΔL6-MD-SNAP was added for 2 mins, then washed twice with T50TC. (4) Finally, T50TC supplemented with 20 mM glucose, 300 μM glucose oxidase, 60 μg/mL catalase, 71 mM ß-mercaptoethanol, and 0.5 μM X-rhodamine MTs was added. For single molecule experiments, the same preparations were performed with the following exceptions. (3) 0.5 μM X-rhodamine/biotin MTs were added for 2 mins, then washed twice with T50TC. (4) as above, except 1-100 nM *Pf*K5ΔL6-MD-SNAP-alexa647 was added instead of MTs.

Fluorescent molecules were visualised on an Eclipse Ti-E inverted microscope with H-TIRF illuminator, LU-N4 laser unit, and CFI Apo TIRF 1.49 N.A. oil objective lens 100x (Nikon) ^60^. Images were recorded on an iXon DU888 Ultra EMCCD camera (Andor) with 60-100 ms exposures. MT velocity and parameters of *Pf*K5ΔL6-MD-SNAP MT binding were calculated from kymographs generated in FIJI ^56^. For MT binding, only events >= 3 frames were considered. Mean dwell time/k_off_ was calculated by fitting a one-phase exponential decay model to the data. Background k_on_ rates were measured by randomly sampling areas without MTs in the ADP and ATP states, because these states had high background binding, owing to the high concentrations needed to observe *Pf*K5ΔL6-MD-SNAP binding events

### Cryo-EM sample preparation, imaging, and data processing

Nucleotide-free *Pf*K5ΔL6-MD was prepared by incubation with 10 U/mL apyrase (Sigma) for at least 15 mins at 4 ^°^C, to remove any nucleotides still present following purification. UltrAuFoil 1.2/1.3® gold (Quantifoil) electron microscopy grids were glow discharged in air for 1 min at 0.3 mpa using a PELCO easiGlow™(Ted Pella, Inc.). Using a Vitrobot Mark IV (ThermoFisher), with the chamber set to 23^°^C and 95 % humidity, 4 μL of 3.5 μM GMPCPP MTs were added to grids and incubated for 30 s, then blotted. 4 μL of 50 μM no nucleotide *Pf*K5ΔL6-MD was then immediately added, incubated for 30 s, then blotted. A second round of *Pf*K5ΔL6-MD was added before blotting and plunge freezing into an ethane slush. AMPPNP *Pf*K5ΔL6-MD was prepared by incubation with 5 mM AMPPNP (R&D Systems) for at least 5 mins at 4 ^°^C. Vitrification was performed as above, except using C-flat 2/2™ carbon (Protochips) holey grids with 5 μM GMPCPP MTs.

Imaging was manually or automatically performed on a G2 Polara (FEI/ThermoFisher) operating at 300 kV using SerialEM ^61^. Images were collected on a K2 summit detector in counting mode, with a GIF Quantum LS Imaging Filter (Gatan). The sample was exposed with 58 e^-^/ Å^2^ for 18 sec and 60 frames collected with a pixel size at the sample of 1.39 Å.

All image processing steps were performed with RELION v3.0 and 3.1 ^27,62^ except where otherwise noted. Beam induced motion in micrographs was corrected using RELION’s implementation of MotionCor2, and the CTF was determined for each micrograph using Gctf ^63^. The start/end coordinates of each MT were manually assigned, and MT particles with a box size of 432 pixels were extracted every 82 Å and normalised. Alignment and asymmetric reconstruction of 14 protofilament (PF) MTs (which are the dominant MT type in GMPCPP preparations ^64^) was performed using MiRP ^28^, as follows, with new MiRP updates noted. RELION parameters used for MiRP PF sorting, initial seam alignment and seam checking steps are listed in supp. table 1. Briefly, PF number assignment for each MT was performed with supervised 3D classification. As part of the MiRP update undertaken during this work, after 3D classification, PF number class assignments for each MT were smoothened by calculating the mode of class assignment over a 7 particle window. Where changes in class assignment occurred within a single MT – due to for example, changes in PF number or major defects within a single MT - MT regions were subsequently treated as distinct MTs. This improved the homogeneity of each MT, increasing confidence in protofilament number assignment and seam location determination. Initial seam alignment was then performed with several iterations of 3D alignment. This was followed, as previously, by Rot angle and X/Y coordinate fitting – however, a local search step was added to improve Rot angle and X/Y shift assignment. Seam checking via supervised 3D classification was then performed, and MTs with less than 50 % confidence in seam class assignment were removed. C1 reconstructions were obtained with a 3D auto-refine run (using the parameters for X/Y refine in supp. table 1, with a solvent mask obtained from a 3D reconstruction of seam checking results), followed by per-particle CTF refinement, Bayesian polishing, and beam-tilt estimation, then a second 3D auto-refine with these new corrections. MiRP was also updated to improve useability, by creating three programs for each MiRP step that can be operated from the RELION v3.1 GUI (https://github.com/moores-lab/MiRPv2).

Symmetrised maps were obtained by first performing 2D classification without alignment (200 classes, T = 8), and selecting well-aligned classes with many particles and an estimated resolution better than 6 Å. A 3D auto-refine run was then performed where 14-fold local symmetry was applied, as previously described ^65^. To address *Pf*K5ΔL6-MD heterogeneity, 3D classification was performed at the level of a *Pf*K5ΔL6-MD:αβ-tubulin asymmetric unit. For this, symmetry expansion was applied to all particles, then 3D classification (3 classes, T = 256) without alignment and a mask around one *Pf*K5ΔL6-MD site opposite the seam was applied. This resulted in one class with clear *Pf*K5ΔL6-MD decoration, which was selected and subjected to a 3D auto-refine procedure. The MT-bound nucleotide-free and AMPPNP-bound *Pf*K5ΔL6-MD reconstructions are deposited with the Electron Microscopy Data Bank, deposition number 12257 and 12258 respectively.

### Sequence analysis, comparative modelling and flexible fitting

To obtain a kinesin-5 family sequence alignment, the motor domains all kinesin-5 family members in the Swiss-Prot database were aligned with MAFFT ^66^. A hidden Markov model of this alignment was created, and queried against the UniProt Reference Proteomes database using HMMER ^67^. Sequences obtained were then compared to a kinesin profile from the Pfam database ^68^, and those with less than 400 identities were removed. Finally, the sequences were aligned with MAFFT, using the L-INS-i method (S12 Data). Secondary structure prediction was performed using Quick2D ^69^, using various prediction algorithms ^70–74^. A residue was assigned helical or beta-sheet identity if 3 or more prediction algorithms agreed.

The above sequence alignment was used in homology modelling of *Pf*K5ΔL6-MD, using the human kinesin-5 AMPPNP bound crystal structure (PDB ID: 3HQD ^34^) as a template. For this, 100 homology models were produced with Modeller v9.2 ^75^, then scored using QMEAN ^76^ and the top model selected. Restraints used to model the extended helixα2 and for highly conserved positions in the nucleotide binding site are listed in supp. table 2. Flexible fitting of *Pf*K5ΔL6-MD secondary structure elements into no nucleotide and AMPPNP cryo-EM reconstructions was performed using Flex-EM ^77^ (cap shift = 0.15) after rigid body docking of the top homology model using the *Fit in Map* tool in Chimera ^78^. During this procedure, the nucleotide binding site (AMPPNP, Mg^2+^, switch-I/II loops, and the P-loop) was defined as a rigid body, while loop regions were treated as “all atoms”. Models of the N-terminus, loops 2, 8, 9, 10, 11, and 12, and the neck linker were predicted using Rosetta, firstly using a coarse method (500 models using cyclic coordinate descent with fragment insertion in the centroid modelling step ^79^), then the model with highest cross-correlation was selected for a second prediction (500 models using kinematic closure with a fit to density term in the centroid modelling step ^80^). A local all-atom fit to density step was then performed using the Rosetta *Relax* procedure including a fit to density term ^81^. Finally, the interface between *Pf*K5ΔL6-MD and αβ-tubulin was refined with protein-protein docking restrained by cryo-EM density in HADDOCK ^82^ as described previously ^31^, using the PDB ID 3JAT ^83^ as αβ-tubulin atomic model. SMOC scores were calculated using the TEMPy software package ^84,85^. The molecular models of MT-bound nucleotide-free and AMPPNP-bound *Pf*K5ΔL6-MD are deposited with the Worldwide Protein Data Bank, deposition number 7NB8 and 7NBA respectively.

### Visualisation and analysis

Plotting was performed with GraphPad Prism 8. Cryo-EM density and model analysis was done in Chimera ^86^ and ChimeraX ^87^. Protein sequence analysis was done in Jalview ^88^. Protein interface areas were calculated with PDBe PISA v1.52 ^89^.

## ACKNOWLEDGEMENTS

A.D.C. was supported by PhD studentships from the Medical Research Council, U.K. (MR/J003867/1) and funds from the Wellcome Trust (101311-10). This work was also supported by funds from the Wellcome Trust (085945/Z/08/Z) to C.A.M., A.J.R (104196/Z/14/Z and 217186/Z/19/Z) and M.T. (209250/Z/17/Z and 208398/Z/17/Z) and the Medical Research Council, U.K. to C.A.M (MR/R000352/1). R.T. was supported by the Biotechnology and Biological Sciences Research Council, U.K. (BB/N017609/1). Cryo-EM data were collected at the Institute of Structural and Molecular Biology (ISMB), Birkbeck on equipment funded by the Wellcome Trust, U.K. (079605/Z/06/Z) and the Biotechnology and Biological Sciences Research Council, U.K. (BB/L014211/1). We thank N. Lukoyanova and S. Chen for electron microscopy support, D. Houldershaw for computational support, A. Peña for the gift of *Hs*K5 protein, member of the Moores and Topf groups for helpful discussions, and C. Hoey, Z. Ahmed and S. Lacey for early work on this project

## SUPPORTING INFORMATION CAPTIONS

**S1 Figure. Biochemical behaviour of *Pf*K5ΔL6-MD constructs**. (A) Effect of pH on *Pf*K5ΔL6-MD ATPase rate, with PIPES used for pH 6.8, HEPES for pH 7.2 and 7.6, and Tris-HCl for Ph 8.0. n = 3 (technical replicates). The mean and 95 % confidence interval are plotted. A constant of 200 μM MTs was used. (B) As in (A), for the effect of KCl concentration on *Pf*K5ΔL6-MD ATPase activity. (C) Example kymographs from *Pf*K5ΔL6-MD-SNAP mediated MT gliding experiments, annotated with measured velocities. (D) Raw data from *Pf*K5ΔL6-MD-SNAP single molecule MT binding experiments, with different nucleotide treatments (NN = no nucleotide). On the left, the MT reference image is shown, and on the right is an example single frame from the corresponding *Pf*K5ΔL6-MD-SNAP movie.

**S2 Figure. Cryo-EM 3D reconstruction and molecule modelling of *Pf*K5ΔL6-MD no nucleotide and AMPPNP states**. (A) Overview of the 3D image processing strategy employed, with reconstructions of the *Pf*K5ΔL6-MD AMPPNP for each step shown, and coloured by local resolution. Firstly, alignment of asymmetric 14 protofilament MTs is achieved with MiRP steps. Then, a high resolution alignment and reconstruction is performed without applying symmetry, and per-particle CTF and motion correction is performed. A symmetrised reconstruction using local symmetry in RELION can then be created, and improved using 2D classification to select for optimal particles. To improve decay in *Pf*K5ΔL6-MD resolution, an asymmetric unit refinement step is performed where symmetry expansion and 3D classification are used to obtain a more homogeneous subset of *Pf*K5ΔL6-MD motors. (B) C1 (unsymmetrised) *Pf*K5ΔL6-MD/microtubule reconstructions for the *Pf*K5ΔL6-MD no nucleotide and AMPPNP states, coloured according to the local resolution scheme in (A). (C) *Pf*K5ΔL6-MD no nucleotide and AMPPNP state structures after asymmetric unit refinement. Surface colouring is according to the local resolution scheme in (A). On the left, the central portion of the *Pf*K5ΔL6-MD/microtubule reconstructions, with *Pf*K5ΔL6-MD enriched at one site. On the right, the *Pf*K5ΔL6-MD/αβ-tubulin asymmetric unit extracted from the microtubule reconstruction on the left.

**S3 Figure. Secondary structure conformations of the *Pf*K5ΔL6-MD no nucleotide and AMPPNP states**. (A) Alignment of the no nucleotide and AMPPNP state cryo-EM maps on αβ-tubulin, showing that conformational changes between the no nucleotide and AMPPNP state can be observed in the cryo-EM maps. (B) The fit to density of the models for various *Pf*K5ΔL6-MD secondary structure elements, showing that α-helices and β-sheets are well resolved. (C) Per-residue TEMPy SMOC scores ^84,85^ for the no nucleotide and AMPPNP state models, that indicate the fit of the model to cryo-EM maps. The SMOC score for the homology model is also show, to demonstrate how the flexible fitting process has improved the models fit to density.

**S4 Figure. *Pf*K5ΔL6-MD AMPPNP state nucleotide density**. To illustrate cryo-EM density corresponding to AMPPNP, synthetic density corresponding to the protein components of the *Pf*K5ΔL6-MD model – i.e. without AMPPNP - was calculated at 6 Å and was subtracted from *Pf*K5ΔL6-MD AMPPNP state cryo-EM reconstruction. The resulting difference density corresponds to the bound nucleotide (modelled as AMPPNP given the sample preparation conditions), αβ-tubulin, and loop5, which was not included in the model.

**S5 Figure. Primary sequence alignment of *Pf*K5 and *Hs*K5**. Coloured by the ClustalX scheme. Secondary structure elements corresponding to the *Pf*K5ΔL6-MD AMPPNP structure are shown, in addition to secondary structure prediction for *Pf*K5.

**S6 Figure. *Pf*K5ΔL6-MD cover-neck bundle formation**. (a) Sequence alignment of helixα6 and the neck-linker, with conservation score from Jalview below. % sequence identity between the different kinesins is noted on the right. (b) View of the *Pf*K5ΔL6-MD no nucleotide and AMPPNP reconstructions showing increased density for the N-terminus in the AMPPNP state, supporting cover neck bundle formation.

**S7 Figure. Sequence conservation between *P. falciparum* and *H. sapiens* αβ-tubulin**. *P. falciparum* α1-tubulin (Uniprot ID: P14642) and β-tubulin (P14643), and *H. sapiens* α-tubulin (Uniprot ID: Q71U36) and β-tubulin (P07437), were aligned in with MAFFT ^66^ using the L-INS-I method, and visualised in Jalview ^88^. Black lines indicate the sections of tubulin that contribute to the kinesin binding site.

**S8 Table. MiRP Alignment parameters**

**S9 Table. Restraints used in *Pf*K5ΔL6-MD homology model generation**

**S10 Table. Comparative rotation of *Pf*K5ΔL6-MD helices between the no nucleotide and AMPPNP states**.

**S11 Table. Residue-residue contacts between the *Pf*K5ΔL6-MD no nucleotide and AMPPNP states, and αβ-tubulin. Residues within contact distance were detected in Chimera**.

**S12 Data. Kinesin-5 motor domain sequence alignment**.

## REFERENCES

1. World Health Organization. World malaria report. (2018).

2. Dondorp, A. M. et al. Artemisinin Resistance in Plasmodium falciparum Malaria. N. Engl. J. Med. 361, 455–467 (2009).

3. World Health Organization. Artemisinin resistance and artemisinin-based combination therapy efficacy. (2018).

4. Birnbaum, J. et al. A Kelch13-defined endocytosis pathway mediates artemisinin resistance in malaria parasites. Science 367, 51–59 (2020).

5. Cowman, A. F., Healer, J., Marapana, D. & Marsh, K. Malaria: Biology and Disease. 167, 610–624 (2016).

6. Van Vuuren, R. J., Visagie, M. H., Theron, A. E. & Joubert, A. M. Antimitotic drugs in the treatment of cancer. 76, 1101–1112 (2015).

7. Hirokawa, N., Noda, Y., Tanaka, Y. & Niwa, S. Kinesin superfamily motor proteins and intracellular transport. 10, 682–96 (2009).

8. Cross, R.a. & McAinsh, A. Prime movers: the mechanochemistry of mitotic kinesins. 15, 257–271 (2014).

9. Wojcik, E. J. Kinesin-5: Cross-bridging mechanism to targeted clinical therapy. 531, 133–149 (2013).

10. von Loeffelholz, O. & Moores, C. A. Cryo-EM structure of the Ustilago maydis kinesin-5 motor domain bound to microtubules. 1–5 (2019).

11. Wickstead, B., Gull, K. & Richards, T. A. Patterns of kinesin evolution reveal a complex ancestral eukaryote with a multifunctional cytoskeleton. 10, 1–12 (2010).

12. Kapitein, L. C. Microtubule cross-linking triggers the directional motility of kinesin-5. 182, 421–428 (2008).

13. Shimamoto, Y., Forth, S. & Kapoor, T. M. Measuring Pushing and Braking Forces Generated by Ensembles of Kinesin-5 Crosslinking Two Microtubules. 34, 669–681 (2015).

14. Acar, S. The bipolar assembly domain of the mitotic motor kinesin-5. 4, 1343 (2013).

15. Myers, S. M. & Collins, I. Recent findings and future directions for interpolar mitotic kinesin inhibitors in cancer therapy. 8, 463–489 (2016).

16. Waitzman, J. S. The loop 5 element structurally and kinetically coordinates dimers of the human kinesin-5, Eg5. 101, 2760–2769 (2011).

17. Larson, A. G., Naber, N., Cooke, R., Pate, E. & Rice, S. E. The conserved L5 loop establishes the pre-powerstroke conformation of the kinesin-5 motor, Eg5. 98, 2619–2627 (2010).

18. Behnke-Parks, W. M. Loop L5 acts as a conformational latch in the mitotic kinesin Eg5. J. Biol. Chem.286, 5242–5253 (2011).

19. Liu, L., Parameswaran, S., Liu, J., Kim, S. & Wojcik, E. J. Loop 5-directed compounds inhibit chimeric Kinesin-5 motors: Implications for conserved allosteric mechanisms. 286, 6201–6210 (2011).

20. Pizzi, E. & Frontali, C. Low-Complexity Regions in Plasmodium falciparum Proteins. 11, 218–229 (2001).

21. Liu, L., Richard, J., Kim, S. & Wojcik, E. J. Small molecule screen for candidate antimalarials targeting Plasmodium Kinesin-5. J. Biol. Chem. 289, 16601–16614 (2014).

22. Peña, A., Sweeney, A., Cook, A. D., Topf, M. & Moores, C. A. Structure of Microtubule-Trapped Human Kinesin-5 and Its Mechanism of Inhibition Revealed Using Cryoelectron Microscopy. Structure 28, 450-457.e5 (2020).

23. Bell, K. M., Cha, H. K., Sindelar, C.V & Cochran, J. C. The yeast kinesin-5 Cin8 interacts with the microtubule in a noncanonical manner. J. Biol. Chem. 292, 14680–14694 (2017).

24. Cochran, J. C. et al. Mechanistic analysis of the mitotic kinesin Eg5. J. Biol. Chem. 279, 38861–38870 (2004).

25. Kapitein, L. C. et al. The bipolar mitotic kinesin Eg5 moves on both microtubules that it crosslinks. Nature 435, 114–118 (2005).

26. Kaseda, K., Crevel, I., Hirose, K. & Cross, R. A. Single-headed mode of kinesin-5. EMBO Rep. 9, 761–765 (2008).

27. Nakane, T. et al. New tools for automated high-resolution cryo-EM structure determination in RELION-3. Elife 7, 1–38 (2018).

28. Cook, A. D., Manka, S. W., Wang, S., Moores, C. A. & Atherton, J. A microtubule RELION-based pipeline for cryo-EM image processing. J. Struct. Biol. 209, p(2020).

29. Manka, S. W. & Moores, C. A. Microtubule structure by cryo-EM: snapshots of dynamic instability. Essays Biochem. 62, 737–751 (2018).

30. Amos, L. A. & Hirose, K. The structure of microtubule-motor complexes. Curr. Opin. Cell Biol. 9, 4–11 (1997).

31. Atherton, J. et al. The divergent mitotic kinesin MKLP2 exhibits atypical structure and mechanochemistry. Elife 6, (2017).

32. Debs, G. E., Cha, M., Liu, X., Huehn, A. R. & Sindelar, C. V. Dynamic and asymmetric fluctuations in the microtubule wall captured by high-resolution cryoelectron microscopy. Proc. Natl. Acad. Sci. U. S. A. 117, 16976–16984 (2020).

33. Vale, R. D. Switches, latches, and amplifiers: Common themes of G proteins and molecular motors. J. Cell Biol. 135, 291–302 (1996).

34. Parke, C. L., Wojcik, E. J., Kim, S. & Worthylake, D. K. ATP hydrolysis in Eg5 kinesin involves a catalytic two-water mechanism. J. Biol. Chem. 285, 5859–5867 (2010).

35. Cao, L. et al. The structure of apo-kinesin bound to tubulin links the nucleotide cycle to movement. Nat. Commun. 5, 1–9 (2014).

36. Hwang, W., Lang, M. J. & Karplus, M. Force Generation in Kinesin Hinges on Cover-Neck Bundle Formation. Structure 16, 62–71 (2008).

37. Rice, S. et al. A structural change in the kinesin motor protein that drives motility. Nature 402, 778–84 (1999).

38. Atherton, J. et al. Conserved mechanisms of microtubule-stimulated ADP release, ATP binding, and force generation in transport kinesins. Elife 3, (2014).

39. von Loeffelholz, O., Peña, A., Drummond, D. R., Cross, R. & Moores, C. A. Cryo-EM Structure (4.5-Å) of Yeast Kinesin-5–Microtubule Complex Reveals a Distinct Binding Footprint and Mechanism of Drug Resistance. J. Mol. Biol. 431, 864–872 (2019).

40. Behnke-Parks, W. M. et al. Loop L5 acts as a conformational latch in the mitotic kinesin Eg5. J. Biol. Chem. 286, 5242–5253 (2011).

41. Kaan, H. Y. K., Ulaganathan, V., Hackney, D. D. & Kozielski, F. An allosteric transition trapped in an intermediate state of a new kinesin-inhibitor complex. Biochem. J. 425, 55–60 (2010).

42. DeBonis, S. et al. In vitro screening for inhibitors of the human mitotic kinesin Eg5 with antimitotic and antitumor activities. Mol. Cancer Ther. 3, 1079–90 (2004).

43. Cahu, J. & Surrey, T. Motile microtubule crosslinkers require distinct dynamic properties for correct functioning during spindle organization in Xenopus egg extract. J. Cell Sci. 122, 1295–1300 (2009).

44. Dowling, D. P. et al. Crystal structure of arginase from plasmodium falciparum and implications for l -arginine depletion in malarial infection. Biochemistry 49, 5600– 5608 (2010).

45. Davies, H. M., Nofal, S. D., McLaughlin, E. J. & Osborne, A. R. Repetitive sequences in malaria parasite proteins. FEMS Microbiol. Rev. 41, 923–940 (2017).

46. Shang, Z. et al. High-resolution structures of kinesin on microtubules provide a basis for nucleotide-gated force-generation. Elife 3, 1–27 (2014).

47. Goulet, A. et al. Comprehensive structural model of the mechanochemical cycle of a mitotic motor highlights molecular adaptations in the kinesin family. 111, 1837–42 (2014).

48. Rai, A. Taxanes convert regions of perturbed microtubule growth into rescue sites. 19, 355–365 (2020).

49. Muretta, J. M. Loop L5 assumes three distinct orientations during the ATPase cycle of the mitotic kinesin Eg5: A transient and time-resolved fluorescence study. 288, 34839–34849 (2013).

50. Yan, Y. Inhibition of a Mitotic Motor Protein: Where, How, and Conformational Consequences. J. Mol. Biol. 335, 547–554 (2004).

51. Zeeshan, M. et al. Plasmodium berghei Kinesin-5 Associates With the Spindle Apparatus During Cell Division and Is Important for Efficient Production of Infectious Sporozoites. Front. Cell. Infect. Microbiol. 10, 1–14 (2020).

52. Zhang, A. M. et al. Uncovering the essential genome of the human malaria parasite Plasmodium falciparum by saturation mutagenesis. Science 360, 1–10 (2018).

53. Yahiya, S., Rueda-Zubiaurre, A., Delves, M. J., Fuchter, M. J. & Baum, J. The antimalarial screening landscape—looking beyond the asexual blood stage. Curr. Opin. Chem. Biol. 50, 1–9 (2019).

54. Aleshnick, M., Ganusov, V. V., Nasir, G., Yenokyan, G. & Sinnis, P. Experimental determination of the force of malaria infection reveals a non-linear relationship to mosquito sporozoite loads. PLoS Pathog. 16, 1–23 (2020).

55. Chua, P. R. et al. Effective killing of the human pathogen Candida albicans by a specific inhibitor of non-essential mitotic kinesin Kip1p. Mol. Microbiol. 65, 347–362 (2007).

56. Schindelin, J. et al. Fiji: An open-source platform for biological-image analysis. Nat. Methods 9, 676–682 (2012).

57. Gibson, D. G. et al. Enzymatic assembly of DNA molecules up to several hundred kilobases. Nat. Methods 6, 343–5 (2009).

58. Hyman, A. A. Preparation of marked microtubules for the assay of the polarity of microtubule-based motors by fluorescence. in Journal of Cell Science 14, 125–127 (1991).

59. Kreuzer, K. N. & Jongeneel, C. V. Escherichia coli Phage T4 Topoisomerase. Methods Enzymol. 100, 144–160 (1983).

60. Toropova, K., Mladenov, M. & Roberts, A. J. Intraflagellar transport dynein is autoinhibited by trapping of its mechanical and track-binding elements. Nat. Struct. Mol. Biol. 24, 461–468 (2017).

61. Mastronarde, D. N. Automated electron microscope tomography using robust prediction of specimen movements. J. Struct. Biol. 152, 36–51 (2005).

62. Zivanov, J., Nakane, T. & Scheres, S. H. W. Estimation of high-order aberrations and anisotropic magnification from cryo-EM data sets in RELION-3.1. IUCrJ 7, 253–267 (2020).

63. Zhang, K. Gctf: Real-time CTF determination and correction. J. Struct. Biol. 193, 1–12 (2016).

64. Vale, R. D., Coppin, C. M., Malik, F., Kull, F. J. & Milligan, R. A. Tubulin GTP hydrolysis influences the structure, mechanical properties, and kinesin-driven transport of microtubules. J. Biol. Chem. 269, 23769–23775 (1994).

65. Lacey, S. E., He, S., Scheres, S. H. W. & Carter, A. P. Cryo-EM of dynein microtubule-binding domains shows how an axonemal dynein distorts the microtubule. Elife 8, 1– 21 (2019).

66. Katoh, K. & Standley, D. M. MAFFT multiple sequence alignment software version 7: improvements in performance and usability. Mol. Biol. Evol. 30, 772–80 (2013).

67. Eddy, S. R. Accelerated profile HMM searches. PLoS Comput. Biol. 7, 1–16 (2011).

68. Bateman, A. The Pfam protein families database. Nucleic Acids Res. 32, 138D – 141 (2004).

69. Zimmermann, L. et al. A Completely Reimplemented MPI Bioinformatics Toolkit with a New HHpred Server at its Core. J. Mol. Biol. 430, 2237–2243 (2018).

70. Jones, D. T. Protein Secondary Structure Prediction Based on Position-specific Scoring Matrices |Elsevier Enhanced Reader. J. Mol. Biol. 292, 195–202 (1999).

71. Heffernan, R., Yang, Y., Paliwal, K. & Zhou, Y. Capturing non-local interactions by long short-term memory bidirectional recurrent neural networks for improving prediction of protein secondary structure, backbone angles, contact numbers and solvent accessibility. Bioinformatics 33, 2842–2849 (2017).

72. Yan, R., Xu, D., Yang, J., Walker, S. & Zhang, Y. A comparative assessment and analysis of 20 representative sequence alignment methods for protein structure prediction. Sci. Rep. 3, (2013).

73. Wang, S., Peng, J., Ma, J. & Xu, J. Protein Secondary Structure Prediction Using Deep Convolutional Neural Fields. Sci. Rep. 6, p(2015).

74. Klausen, M. S. et al. NetSurfP-2.0: Improved prediction of protein structural features by integrated deep learning. Proteins Struct. Funct. Bioinforma. 87, 520–527 (2019).

75. Šali, A. & Blundell, T. L. Comparative Protein Modelling by Satisfaction of Spatial Restraints. J. Mol. Biol. 234, 779–815 (1993).

76. Benkert, P., Kunzli, M. & Schwede, T. QMEAN server for protein model quality estimation. Nucleic Acids Res. 37, 1–5 (2009).

77. Topf, M. et al. Protein Structure Fitting and Refinement Guided by Cryo-EM Density. Structure 16, 295–307 (2008).

78. Goddard, T. D., Huang, C. C. & Ferrin, T. E. Visualizing density maps with UCSF Chimera. J. Struct. Biol. 157, 281–287 (2007).

79. Wang, C., Bradley, P. & Baker, D. Protein-Protein Docking with Backbone Flexibility. J. Mol. Biol. 373, 503–519 (2007).

80. Mandell, D. J., Coutsias, E. A. & Kortemme, T. Sub-angstrom accuracy in protein loop reconstruction by robotics-inspired conformational sampling. Nat. Methods 6, 551– 552 (2009).

81. Wang, R. Y. R. et al. Automated structure refinement of macromolecular assemblies from cryo-EM maps using Rosetta. Elife 5, 1–22 (2016).

82. Zundert, G. C. P. Van, Rodrigues, J. P. G. L. M., Trellet, M. & Schmitz, C. The HADDOCK2. 2 Web Server?: User-Friendly Integrative Modeling of Biomolecular Complexes. J. Mol. Biol. 428, 720–725 (2016).

83. Zhang, R., Alushin, G. M., Brown, A. & Nogales, E. Mechanistic origin of microtubule dynamic instability and its modulation by EB proteins. Cell 162, 849–859 (2015).

84. Farabella, I. et al. TEMPy?: a Python library for assessment of three-dimensional electron microscopy density fits. J. Appl. Crystallogr. 48, 1314–1323 (2015).

85. Joseph, A. P. et al. Refinement of atomic models in high resolution EM reconstructions using Flex-EM and local assessment. Methods 100, 42–49 (2016).

86. Pettersen, E. F. et al. UCSF Chimera — A Visualization System for Exploratory Research and Analysis. J. Comput. Chem. 25, 1605–1612 (2004).

87. Goddard, T. D. et al. UCSF ChimeraX: Meeting modern challenges in visualization and analysis. Protein Sci. 27, 14–25 (2018).

88. Waterhouse, A. M., Procter, J. B., Martin, D. M. A., Clamp, M. & Barton, G. J. Jalview Version 2--a multiple sequence alignment editor and analysis workbench. Bioinformatics 25, 1189–1191 (2009).

89. Krissinel, E. & Henrick, K. Inference of Macromolecular Assemblies from Crystalline State. J. Mol. Biol. 372, 774–797 (2007).

