## Supporting Information for "Molecular mechanism of a parasite kinesin motor and implications for its inhibition"

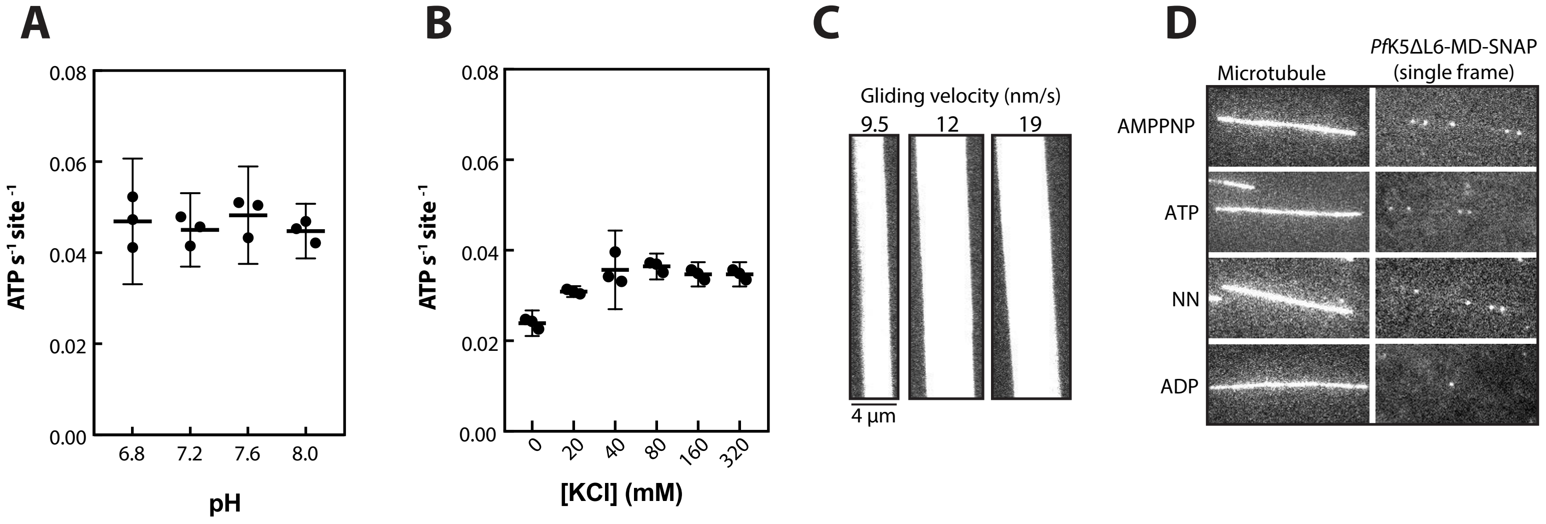

A
0.08
0.06
0.04
0.02
0.00
ATP s-1 site-1
6.8
7.2
7.6
8.0
pH
B
[KCl] (mM)
0
20
40
80
160
320
C
Gliding velocity (nm/s)
9.5
12
19
4 µm
D
Microtubule
PfK5ΔL6-MD-SNAP (single frame)
AMPPNP
ATP
NN
ADP


S1 Fig

S2 Fig

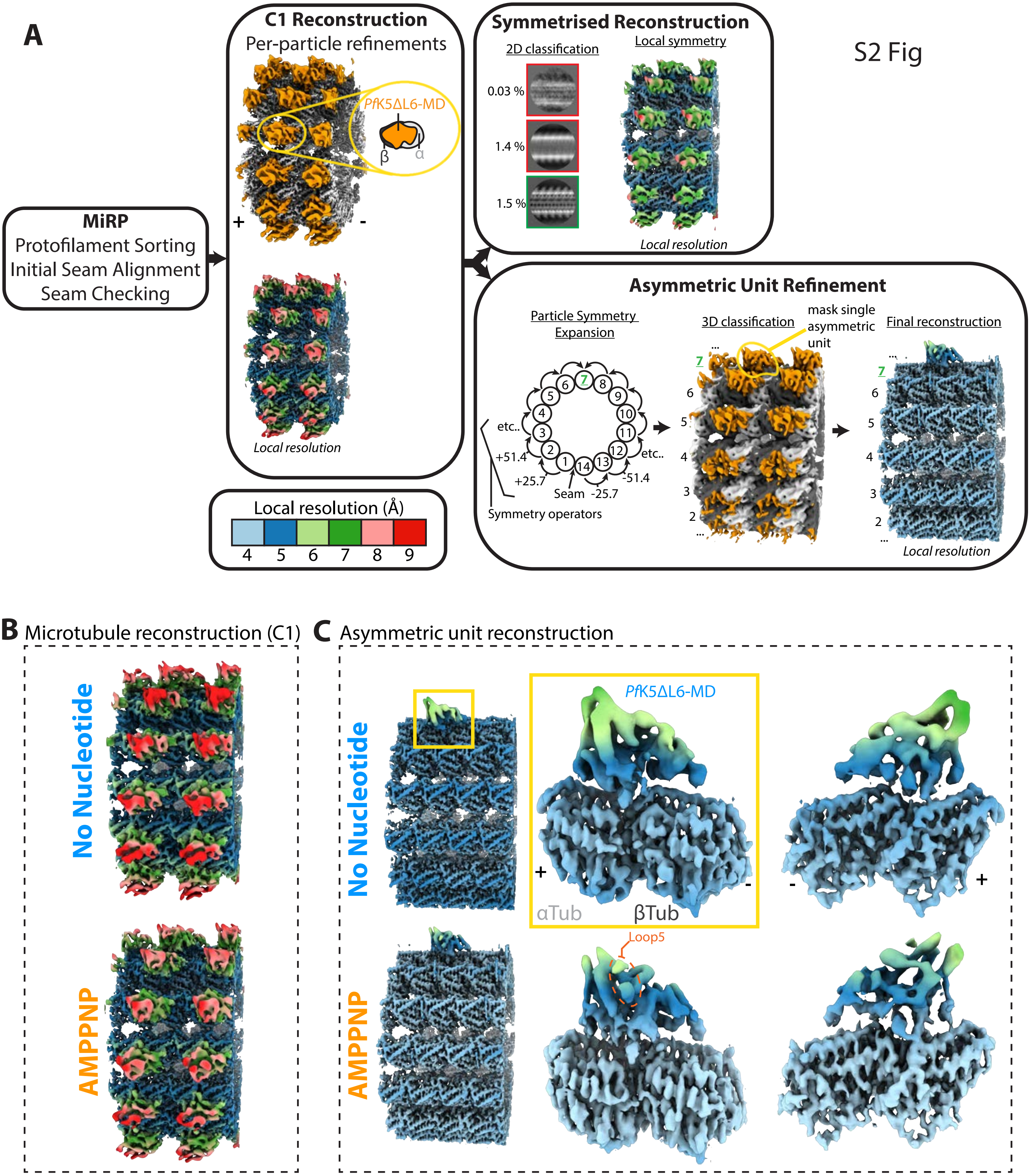

A
MiRP
Protofilament Sorting
Initial Seam Alignment
Seam Checking
C1 Reconstruction
Per-particle refinements
PfK5ΔL6-MD
β
α
+
-
Local resolution
Local resolution (Å)
4 5 6 7 8 9
Symmetrised Reconstruction
2D classification
Local symmetry
0.03 %
1.4 %
1.5 %
Local resolution
Asymmetric Unit Refinement
Particle Symmetry Expansion
3D classification
mask single asymmetric unit
Final reconstruction
etc..
+51.4
+25.7
Seam -25.7
-51.4
etc..
Symmetry operators
Local resolution
B Microtubule reconstruction (C1)
No Nucleotide
AMPPNP
C Asymmetric unit reconstruction
No Nucleotide
PfK5ΔL6-MD
αTub
βTub
Loop5
AMPPNP

S3 Fig

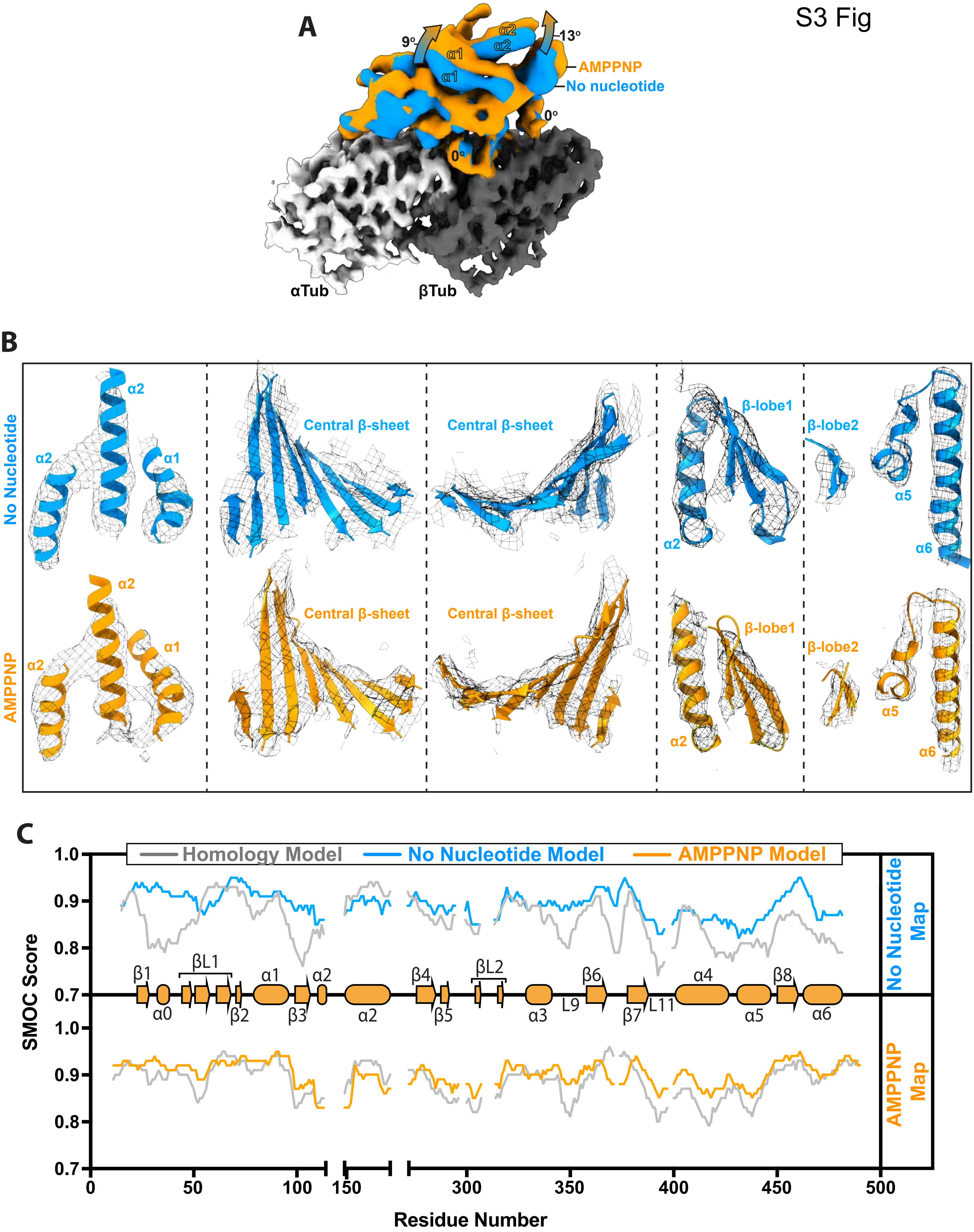
A
9°
13°
α1
α2
AMPPNP
No nucleotide
0°
αTub
βTub
B
No Nucleotide
AMPPNP
Central β-sheet
β-lobe1
β-lobe2
α5
α6
C
Homology Model
No Nucleotide Model
AMPPNP Model
SMOC Score
Residue Number
No Nucleotide Map
AMPPNP Map
β1
βL1
βL2
β2
β3
β4
β5
β6
β7
β8
α0
α3
α4
L9
L11

S4 Fig

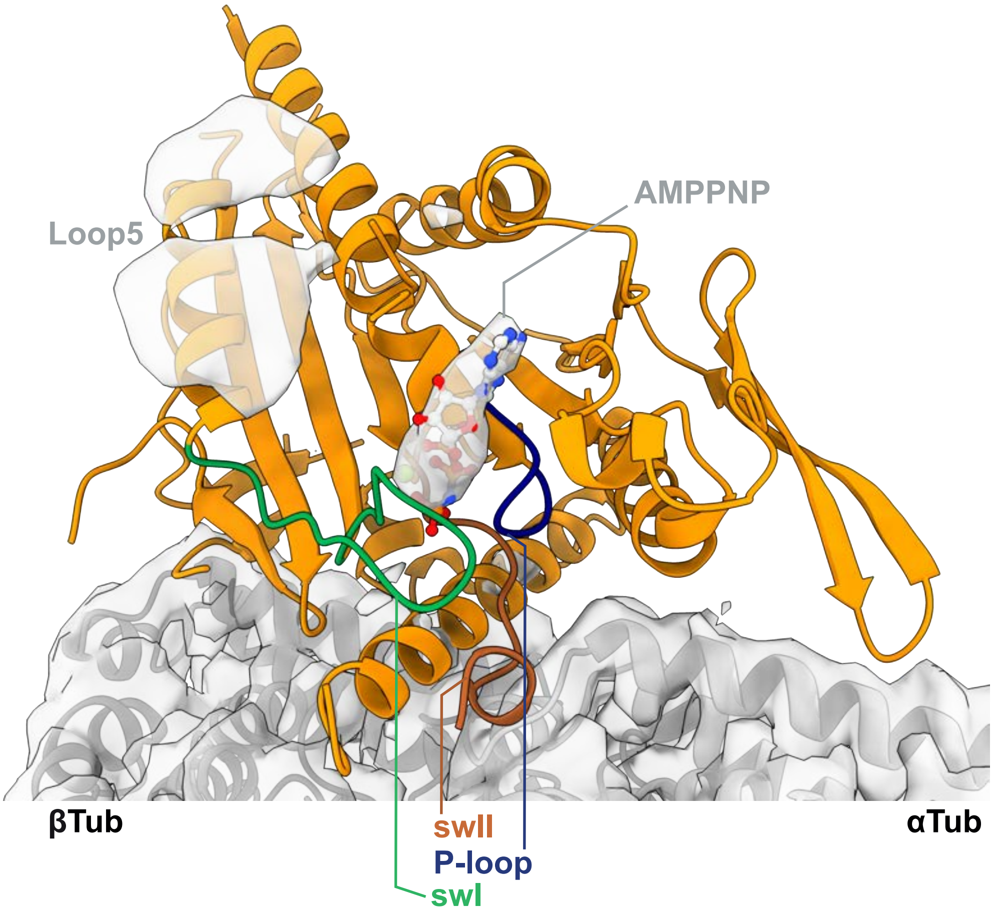

S5 Fig

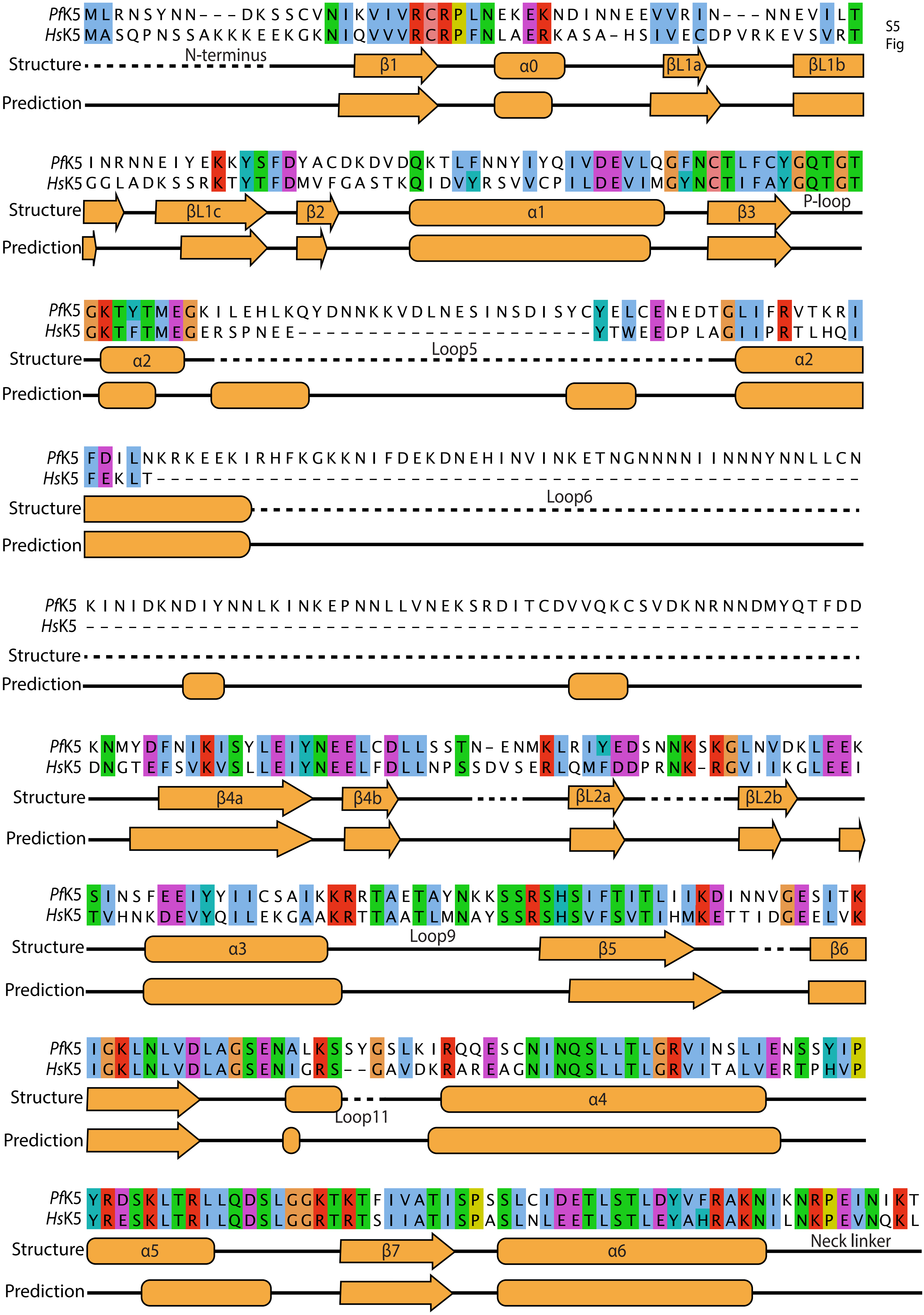
PfK5 MLRNSYNN---DKSSCVNIKVIVRCRPLNEKEKNDINNEEVVRIN---NNEVILT
HsK5 MASQPNSSAKKKEEKGKNIQVVVRCRPFNLAERKASA-HSIVECDPVRKEVSVRT
Structure
N-terminus
β1
α0
βL1a
βL1b
Prediction
PfK5 INRNNEIYEKKYSFDYACDKDVDQKTLFNNYIYQIVDEVLQGFNCTLFCYGQTGT
HsK5 GGLADKSSRKTYTFDMVFGASTKQIDVYRSVVCPILDEVIMGYNCTIFAYGQTGT
Structure
βL1c
β2
α1
β3
P-loop
Prediction
PfK5 GKTYTMEGKILEHLKQYDNNKKVDLNESINSDISYCYELCENEDTGLIFRVTKRI
HsK5 GKTFTMEGERSPNEE---------------------YTWEEDPLAGIIPRTLHQI
Structure
α2
Loop5
α2
Prediction
PfK5 FDILNKRKEEKIRHFKGKKNIFDEKDNEHINVINKETNGNNNNIINNNYNNLLCN
HsK5 FEKLT--------------------------------------------------
Structure
Loop6
Prediction
PfK5 KINIDKNDIYNNLKINKEPNNLLVNEKSRDITCDVVQKCSVDKNRNNDMYQTFDD
HsK5 -------------------------------------------------------
Structure
Prediction
PfK5 KNMYDFNIKISYLEIYNEELCDLLSSTN-ENMKLRIYEDSNNKSKGLNVDKLEEK
HsK5 DNGTEFSVKVSLLEIYNEELFDLLNPSSDVSERLQMFDDPRNK-RGVIIKGLEEI
Structure
β4a
β4b
βL2a
βL2b
Prediction
PfK5 SINSFEEIYYIICSAIKKRRTAETAYNKKSSRSHSIFTITLIIKDINNVGESITK
HsK5 TVHNKDEVYQILEKGAAKRTTAATLMNAYSSRSHSVFSVTIHMKETTIDGEELVK
Structure
α3
Loop9
β5
β6
Prediction
PfK5 IGKLNLVDLAGSENALKSSYGSLKIRQQESCNINQSLLTLGRVINSLIENSSYIP
HsK5 IGKLNLVDLAGSENIGRS--GAVDKRAREAGNINQSLLTLGRVITALVERTPHVP
Structure
α4
Loop11
Prediction
PfK5 YRDSKLTRLLQDSLGGKTKTFIVATISPSSLCIDETLSTLDYVFRAKNIKNRPEINIKT
HsK5 YRESKLTRILQDSLGGRTRTSIIATISPASLNLEETLSTLEYAHRAKNILNKPEVNQKL
Structure
α5
β7
α6
Neck linker
Prediction

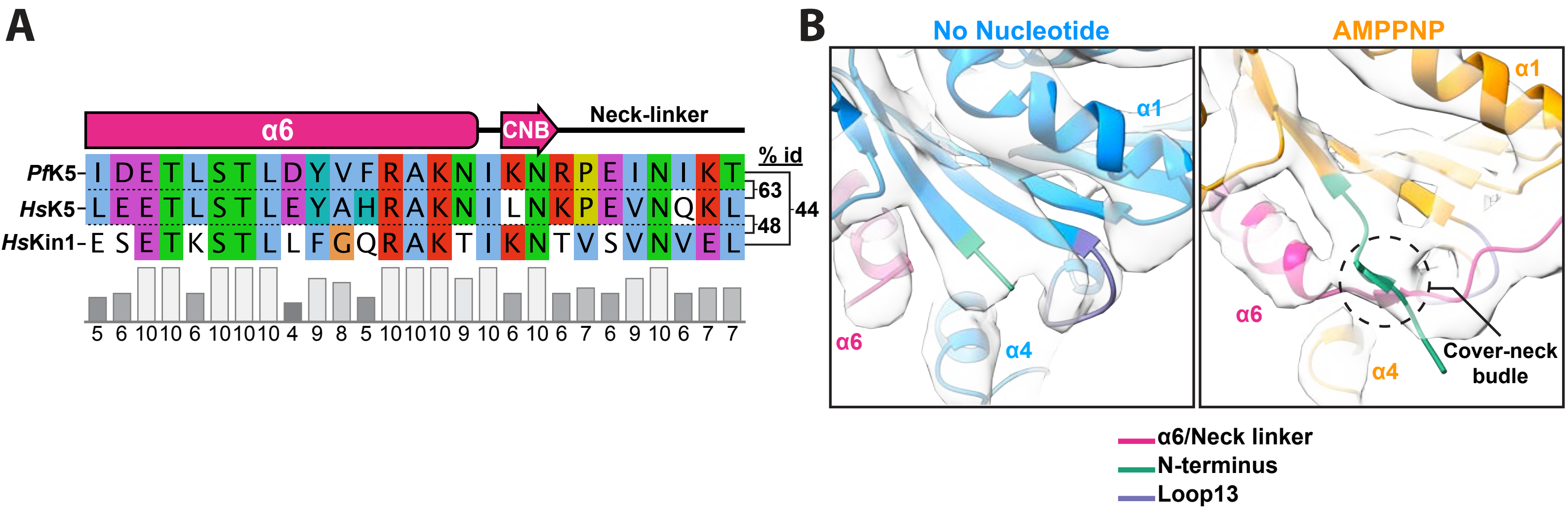
A
α6
CNB
Neck-linker
% id
PfK5
HsK5
HsKin1
I D E T L S T L D Y V F R A K N I K N R P E I N I K T
L E E T L S T L E Y A H R A K N I L N K P E V N Q K L
E S E T K S T L L F G Q R A K T I K N T V S V N V E L
63
48
44
5 6 10 10 6 10 10 10 4 9 8 5 10 10 10 9 10 6 10 6 7 6 9 10 6 7 7
B
No Nucleotide
AMPPNP
α1
α6
α4
Cover-neck budle
α6/Neck linker
N-terminus
Loop13

S6 Fig

### S7 Fig

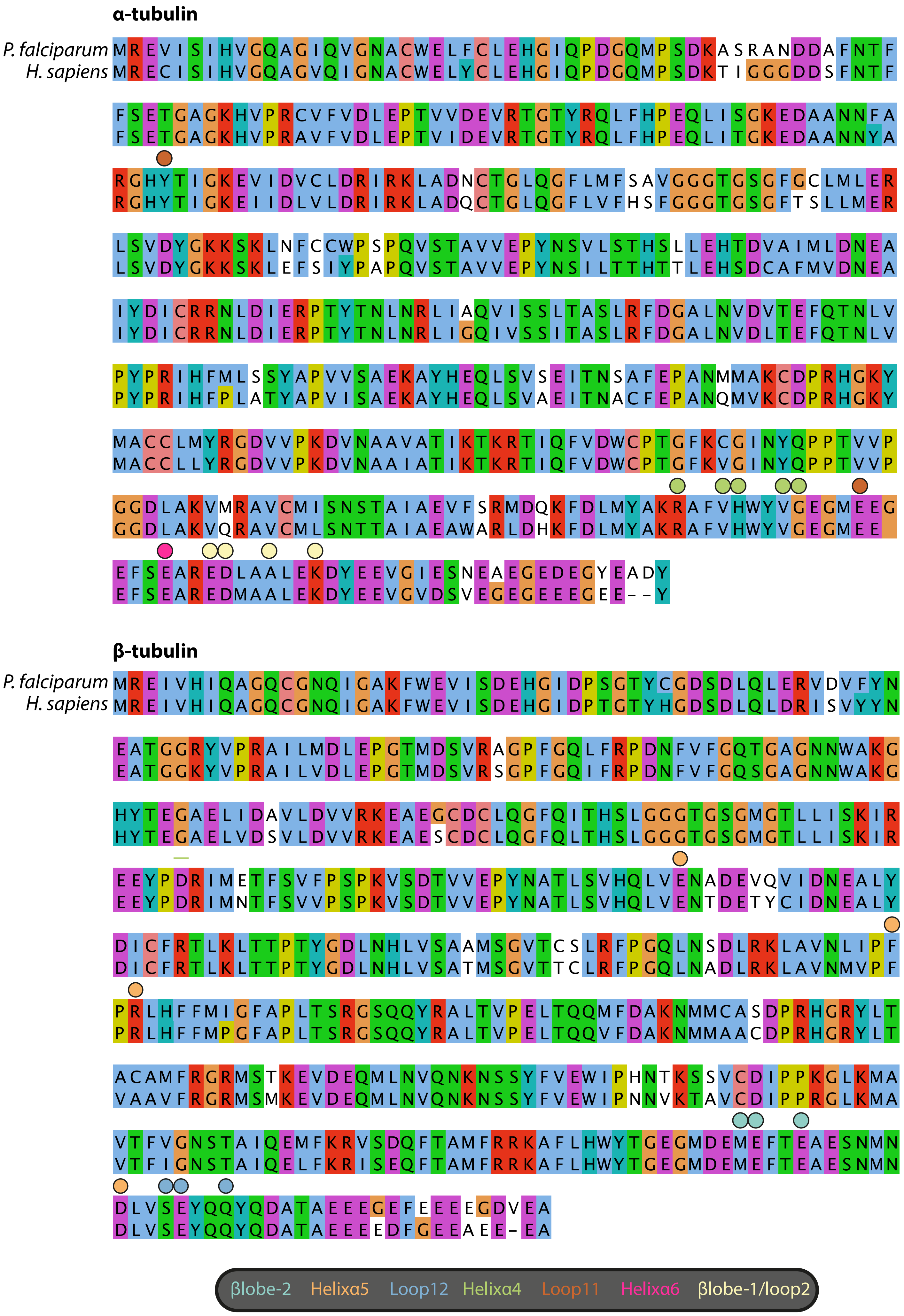
α-tubulin
P. falciparum MREVISIHVGQAGIQVGNACWELFCLEHGIQPDGQMPSDKASRANDDAFNTF
H. sapiens MRECISIHVGQAGVQIGNACWELYCLEHGIQPDGQMPSDKTIGGGDDSFNTF
FSETGAGKHVPRCVFVDLEPTVVDEVRTGTYRQLFHPEQLISGKEDAANNFA
FSETGAGKHVPRAVFVDLEPTVIDEVRTGTYRQLFHPEQLITGKEDAANNYA
RGHYTIGKEVIDVCLDRIRKLADNCTGLQGFLMFSAVGGGTGSGFGCLMLER
RGHYTIGKEIIDLVLDRIRKLADQCTGLQGFLVFHSFGGGTGSGFTSLLMER
LSVDYGKKSKLNFCCWPSPQVSTAVVEPYNSVLSTHSLLEHTDVAIMLDNEA
LSVDYGKKSKLEFSIYPAPQVSTAVVEPYNSILTTHTTLEHSDCAFMVDNEA
IYDICRRNLDIERPTYTNLNRLIAQVISSLTASLRFDGALNVDVTEFQTNLV
IYDICRRNLDIERPTYTNLNRLIGQIVSSITASLRFDGALNVDLTEFQTNLV
PYPRIHFMLSSYAPVVSAEKAYHEQLSVSEITNSAFEPANMMAKCDPRHGKY
PYPRIHFPLATYAPVISAEKAYHEQLSVAEITNACFEPANQMVKCDPRHGKY
MACCLMYRGDVVPKDVNAAVATIKTKRTIQFVDWCPTGFKCGINYQPPTVVP
MACCLLYRGDVVPKDVNAAIATIKTKRTIQFVDWCPTGFKVGINYQPPTVVP
GGDLAKVMRAVCMISNSTAIAEVFSRMDQKFDLMYAKRAFVHWYVGEGMEEG
GGDLAKVQRAVCMLSNTTAIAEAWARLDHKFDLMYAKRAFVHWYVGEGMEE
EFSEAREDLAALEKDYEEVGIESNEAEGEDEGYEADY
EFSEAREDMAALEKDYEEVGVDSVEGEGEEEGEE--Y
β-tubulin
P. falciparum MREIVHIQAGQCGNQIGAKFWEVISDEHGIDPSGTYCGDSDLQLERVDVFYN
H. sapiens MREIVHIQAGQCGNQIGAKFWEVISDEHGIDPTGTYHGDSDLQLDRISVYYN
EATGGRYVPRAILMDLEPGTMDSVRAGPFGQLFRPDNFVFGQTGAGNNWAKG
EATGGKYVPRAILVDLEPGTMDSVRSGPFGQIFRPDNFVFGQSGAGNNWAKG
HYTEGAELIDAVLDVVRKEAEGCDCLQGFQITHSLGGGTGSGMGTLLISKIR
HYTEGAELVDSVLDVVRKEAESCDCLQGFQLTHSLGGGTGSGMGTLLISKIR
EEYPDRIMETFSVFPSPKVSDTVVEPYNATLSVHQLVENADEVQVIDNEALY
EEYPDRIMNTFSVVPSPKVSDTVVEPYNATLSVHQLVENTDETYCIDNEALY
DICFRTLKLTTPTYGDLNHLVSAAMSGVTCSLRFPGQLNSDLRKLAVNLIPF
DICFRTLKLTTPTYGDLNHLVSATMSGVTTCLRFPGQLNADLRKLAVNMVPF
PRLHFFMIGFAPLTSRGSQQYRALTVPELTQQMFDAKNMMCASDPRHGRYLT
PRLHFFMPGFAPLTSRGSQQYRALTVPELTQQVFDAKNMMAACDPRHGRYLT
ACAMFRGRMSTKEVDEQMLNVQNKNSSYFVEWIPHNTKSSVCDIPPKGLKMA
VAAVFRGRMSMKEVDEQMLNVQNKNSSYFVEWIPNNVKTAVCDIPPRGLKMA
VTFVGNSTAIQEMFKRVSDQFTAMFRRKAFLHWYTGEGMDEMEFTEAESNMN
VTFIGNSTAIQELFKRISEQFTAMFRRKAFLHWYTGEGMDEMEFTEAESNMN
DLVSEYQQYQDATAEEEGEFEEEEGDVEA
DLVSEYQQYQDATAEEEEDFGEEAEE-EA
βlobe-2
Helixα5
Loop12
Helixα4
Loop11
Helixα6
βlobe-1/loop2

| Parameter | PF Sorting | Initial Seam Alignment | | | | Seam Check |
|---|---|---|---|---|---|---|
| | | Global Search | Local Search | Rot Refine | X/Y Refine | |
| Offset range | 15 | 15 | 15 | 15 | 4 | – |
| Offset step | 1 | 1 | 1 | 1 | 0.5 | |
| Angular sampling | 1.8 | 0.9 | 0.9 | 0.9 | 0.9 | |
| Sigma Rot | 0 | 0 | 18 | 3 | 3 | |
| Iterations | 1 | 1 | 1 | 10 | 10 | 1 |
| MiRP pre-run | _ | Reset R, X, Y* | Reset X, Y | Reset X, Y | _ | _ |
| MiRP post-run | Vote on PF Number | Vote on Rot | Vote on Rot | Vote on X/Y shifts | – | Vote on seam position |

**S8 Table: MiRP Alignment parameters**

| Start | End | Restraint |
|---|---|---|
| 151 | 172 | α-helix |
| O1B (AMPPNP) | NZ Lys106 | distance (2.7 Å) |
| O1G (AMPPNP) | NZ Lys106 | distance (2.4 Å) |
| CA | NZ Lys106 | distance (5.8 Å) |
| OE1 Glu391 | NH2 Arg355 | distance (2.5 Å) |
| OE2 Glu391 | NH1 Arg355 | distance (4 Å) |
| OD1 Asn412 | N Glu391 | distance (2.9 Å) |
| ND2 Asn412 | O Glu391 | distance (2.8 Å) |

**S9 Table: Restraints used in *Pf*K5ΔL6-MD homology model generation**

| Kinesin motor domain secondary structure elements | Subdomain | Change In angle between no nucleotide and ATP-like states (°) |
|---|---|---|
| α0 | P-loop | 9 |
| α1 | P-loop | 9 |
| α2 | switch-I/II | 13 |
| α3 | switch-I/II | 15 |
| α4 | MT binding | 1 |
| α5 | MT binding | 0 |
| α6 | P-loop | 13 |

**S10 Table: Comparative rotation of *Pf*K5ΔL6-MD helices between the no nucleotide and AMPPNP states.**

| α-tubulin residue | β-tubulin residue | *Pf*K5ΔL6-MD (no nucleotide) | *Pf*K5ΔL6-MD (AMPPNP) | *Pf*K5ΔL6-MD secondary structure element |
|---|---|---|---|---|
| Glu423 | | Arg52 | Arg52 | βlobe-1/loop2 |
| Asp424 | | | Asn53 | |
| Lys430 | | | Glu55 | |
| Glu423 | | Lys59 | Lys59 | |
| | Met406 | | Arg303 | βLobe-2 |
| | Glu407 | | Arg303 | |
| | Glu410 | | Arg303 | |
| | Met406 | | Tyr305 | |
| | Glu410 | Tyr305 | Tyr305 | |
| | Met406 | Glu306 | | |
| | Glu410 | Glu306 | | |
| Glu414 | | | Ser390 | Loop11 |
| Glu414 | | Asn392 | | |
| Tyr108 | | | Leu394 | |
| | Asp161 | Lys402 | Lys402 | Helixα4/loop12 |
| Gly410 | | Gln405 | | |
| Gly410 | | | Cys409 | |
| Val409 | | Asn412 | Asn412 | |
| Gly410 | | Gln413 | Gln413 | |
| Val405 | | Leu416 | | |
| Val409 | | Leu416 | | |
| His406 | | Leu416 | Leu416 | |
| Glu415 | | | Leu416 | |
| Lys401 | | Glu427 | | |
| | Gln424 | | Ser430 | |
| | Asp417 | Tyr431 | Tyr431 | |
| | Ser420 | | Tyr431 | |
| | Glu421 | | Tyr431 | |
| | Gln424 | | Tyr431 | |
| | Glu421 | Ile432 | | Helixα5 |
| | Asp417 | Arg435 | Arg435 | |
| | Arg262 | Arg435 | | |
| | Phe260 | | Asp436 | |
| | Arg262 | Asp436 | Asp436 | |
| | Glu194 | | Lys438 | |
| | Glu410 | Arg441 | | |
| | Asp417 | | Arg441 | |
| Glu420 | | | Asp467 | Helixα5 |
| Glu414 | | Ser471 | | |
| Ser419 | | Asp474 | | |
| Tyr399 | | Arg478 | | |
| Arg402 | | Arg478 | | |
| Ser419 | | Arg478 | | |

**S11 Table: Residue-residue contacts between the *Pf*K5ΔL6-MD no nucleotide and AMPPNP states, and αβ-tubulin. Residues within contact distance were detected in Chimera.**
